# Optimization of lag phase shapes the evolution of a bacterial enzyme

**DOI:** 10.1101/088013

**Authors:** Bharat V. Adkar, Michael Manhart, Sanchari Bhattacharyya, Jian Tian, Michael Musharbash, Eugene I. Shakhnovich

## Abstract

Mutations provide the variation that drives evolution, yet their effects on fitness remain poorly understood. Here we explore how mutations in the essential enzyme Adenylate Kinase (Adk) of *E. coli* affect multiple phases of population growth. We introduce a biophysical fitness landscape for these phases, showing how they depend on molecular and cellular properties of Adk. We find that Adk catalytic capacity in the cell (product of activity and abundance) is the major determinant of mutational fitness effects. We show that bacterial lag times are at a well-defined optimum with respect to Adk’s catalytic capacity, while exponential growth rates are only weakly affected by variation in Adk. Direct pairwise competitions between strains show how environmental conditions modulate the outcome of a competition where growth rates and lag times have a tradeoff, altogether shedding light on the multidimensional nature of fitness and its importance in the evolutionary optimization of enzymes.

## Introduction

Random mutagenesis is often used to assess the distribution of fitness effects in simple experimental models such as propagating viruses and microbes evolving under antibiotic stress^1, 2^. However, the enormous size of sequence space severely constrains how much of the fitness landscape can be explored this way, and mechanistic and predictive insights from these experiments are further limited by a lack of knowledge of the molecular effects of mutations. Instead, a more targeted experimental approach relies on the concept of a biophysical fitness landscape, in which fitness effects of mutations are mapped through their effects on molecular traits of the mutated proteins. In this approach, biophysically-rational genetic variation is introduced on the chromosome, and the molecular and phenotypic effects of that variation are analyzed concurrently^3–6^. By mapping fitness effects to variation of molecular properties rather than to sequences of mutated proteins, we can dramatically reduce the dimensionality of the genotype-to-phenotype mapping. The underlying hypothesis is that variation in a small number of properly-selected molecular traits of mutated proteins can explain most of the resulting mutational variation in fitness, and that the relationship between these molecular traits and fitness is smooth and continuous. Several recent studies have supported this approach^5–7^.

The relationship between sequence variation and fitness is further confounded by the fact that multiple life-history traits contribute to fitness^8^, and the relative importance of these traits to the long-term evolutionary fate of a mutation may be highly dependent on environmental and ecological conditions. While multicellular organisms are generally described by a large number of traits (e.g., viability at various life phases, mating success, fecundity, etc.), unicellular microorganisms like bacteria and yeast are described by relatively fewer components of fitness, such as the time in lag phase, the exponential growth rate, and the overall yield at saturation. All these phases of growth contribute toward the outcome when in competition for limited resources, and hence determine fitness^3,9^. The relative importance of these different phases of bacterial growth in sculpting the fitness landscape depends on the conditions of growth and competition^10–12^.

Overall, the challenge in quantitatively characterizing the fitness landscape is twofold: Understanding fitness in terms of contributions from different phases of growth, and linking each of these components to molecular and cellular traits. In this work, we address both challenges by introducing biophysically-rational genetic variation in the *adk* locus that encodes the essential *E. coli* enzyme Adenylate Kinase (Adk), and projecting the ensuing variations of fitness effects (phenotypic components like growth rate and lag time) onto the biophysical traits of Adk. We find that a unique combination of molecular and cellular traits of Adk — the product of intracellular abundance and catalytic activity, which we term catalytic capacity — provides a reliable predictor of fitness effects across the full range of phenotypic variation. Furthermore, we find that the length of the lag phase is more sensitive to variation in Adk catalytic capacity than is the exponential growth rate, so that the lag phase of the wild-type *E. coli* appears to be optimal with respect to variation in Adk catalytic capacity.

## Results

### Biophysical properties of Adk mutants

Destabilizing mutations have been shown to cause a drop in intracellular protein abundance, mostly through a decrease in the folded fraction of the protein^3^. Hence in order to sample a broad range of molecular and cellular traits of Adk protein below the wild-type levels, we chose a set of 21 missense mutations at 6 different positions of *adk*. (Table S1 and Fig. 1). We selected residues such that their accessible surface area was less than 10% and they were at least 6 Å away from the catalytically-active sites of Adk, so that mutations at these residues were likely to destabilize the protein^13^. For most mutants, we chose amino acid mutations that appeared only at low frequency in an alignment of 895 homologous sequences of Adk. As intended, the purified mutant proteins were destabilized over a wide range (~17 °C in terms of Tm, and ~7.5 kcal/mol in terms of folding Δ*G*) (Table S1, Figs. 1B, S1, S2). In only one case (L209I) did we change the *E. coli* sequence to the consensus amino acid at that position, and we found it in fact stabilized the protein by ~1 kcal/mol (Table S1). Although most of the Adk mutants were less stable than the wild-type (WT), they nevertheless existed predominantly as monomers in solution (Fig. S3). However, several mutations in one position — V106H, V106N, and V106W — did have significant fractions of proteins present in higher oligomeric forms, in addition to the predominant monomeric species (Fig. S3). These proteins bound 4,4’-Dianilino-1,1’-binaphthyl-5,5’-Disulfonate (Bis-ANS) dye to a higher degree compared to the rest of the mutants (Fig. S4), indicating the presence of possible molten globule states in solution^14^. The proteostat dye that reports on protein aggregation^4,15^ also bound these mutants more strongly compared to others (Fig. S4), clearly indicating a higher fraction of aggregated species. The catalytic efficiency (*k*_*cat*_/*K*_*M*_) of the mutant Adk proteins was distributed broadly with most mutants showing a lower activity than *E. coli* WT (Table S1, Figs. 1C, S5).

**Fig. 1:**
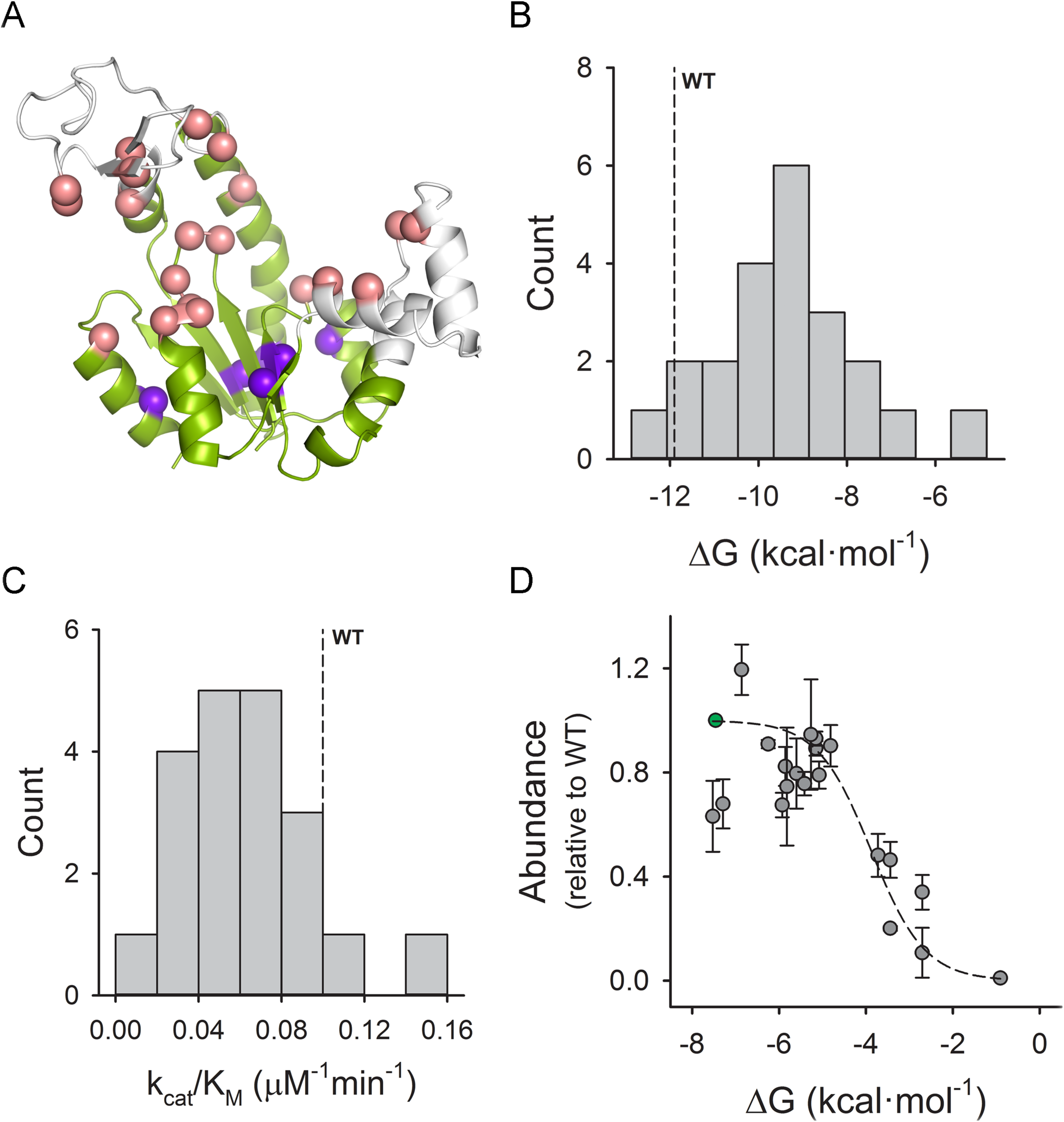
Biophysical and intracellular properties. (A) Crystal structure of Adenylate Kinase from *E. coli* (PDB ID 4ake^30^). The core domain is colored in green, while the LID and NMP domains are shown in white. The C atoms of active-site residues are shown in pink, and the blue spheres represent the C atoms of the 6 buried positions which were mutated in this study. (B) Histogram showing the distribution of folding free energies for all mutant proteins, as determined by isothermal urea denaturation at 25 °C. The stability of WT is marked by a dashed line. (C) Histogram of the catalytic activity parameter *k*_*cat*_/*K*_*M*_ for all mutants. The dashed line indicates the WT value. (D) Total intracellular abundance of mutant Adk proteins as a function of Δ*G* at 37 °C. The abundances are normalized by the WT value. Each data point represents the mean and error bars are standard deviation over two experiments. The dashed line represents the fit to the Boltzmann distribution function described in Eq. 1, where ^*k_B_*^ was 1.987 cal/mol/K. See related Figs. S1-S5 and Table S1.

### Intracellular abundance of Adk follows prediction from Boltzmann distribution

We then incorporated each of the 21 *adk* mutations one-by-one into the *E. coli* chromosome using a genome-editing approach based on homologous recombination^3,4^. We measured the *total* intracellular abundance of WT and mutant Adk proteins using a quantitative western blot (Table S2). The sigmoidal dependence of total intracellular Adk abundance on folding stability Δ*G* (Fig. 1D) is well-described by the Boltzmann distribution for two-state unfolding proteins:

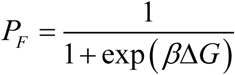

where *P_F_* is the fraction of folded molecules in the ensemble of intracellular Adk and *b* =1/ *k*_*B*_*T*, with Boltzmann constant k_B_ and growth temperature *T*. The total measured abundance of a protein is its amount in the cytoplasm at steady-state, achieved by a balance between production and degradation. Since Adk is expressed from a constitutive promoter in the cells, it is generally safe to assume that the rates of production of all mutants are similar. Under this assumption, the sigmoidal dependence of abundance on stability clearly indicates that the unfolded protein is degraded in the active medium of the cytoplasm.

### Mutations in Adk affect lag times more than exponential growth rates

Mutations in Adk affect both intracellular abundance (via folding stability) and catalytic activity of the protein. Flux dynamics theory predicts, and experiments have confirmed, that the key enzymatic parameter determining the flux through an enzymatic reaction chain is the quantity which we call “catalytic capacity,” defined as the product of intracellular abundance and enzymatic efficiency *k*_*cat*_/*K*_*M*_^5,6,16^. To that end, we determined how two key components of bacterial growth — the exponential growth rate and the lag time (Fig. 2A) — depend on the total catalytic capacity of Adk in *E. coli* cells (Fig. 2B, C; also see *Methods* and Fig. S6-S8 for estimation of growth parameters). We find that while only 3 out of 21 strains show a drop in growth rate greater than 5% of WT, 17 strains show an increase in lag time for a similar change over the WT value (Table S2). This suggests that the mutations in Adk affect the lag phase more significantly than the exponential growth phase. One mechanism for producing longer apparent lag times is when a greater proportion of cells that come out of stationary phase are simply nonviable, as described in a recent study^17^. However, this appears not to be the major cause in our case, as lag times are fairly consistent across replicates (error bars in Fig. 2C) and do not negatively correlate with the number of viable cells (Fig. S9). We also find that the variation in total catalytic capacity of Adk correlates better with the variation in lag times (Spearman’s rank correlation *ρ* = −0.44, *p* = 0.057) than with the variation in growth rates (Spearman’s rank correlation *ρ* = −0.08, *p* = 0.737) (Fig. S10). The variation in lag times is also better explained by the variation in catalytic capacity than with any of the Adk properties separately (stability, abundance, or activity) (Fig. S10). Surprisingly, growth rate appears to tolerate a rather large drop in catalytic capacity of Adk, while lag time does not.

**Fig. 2:**
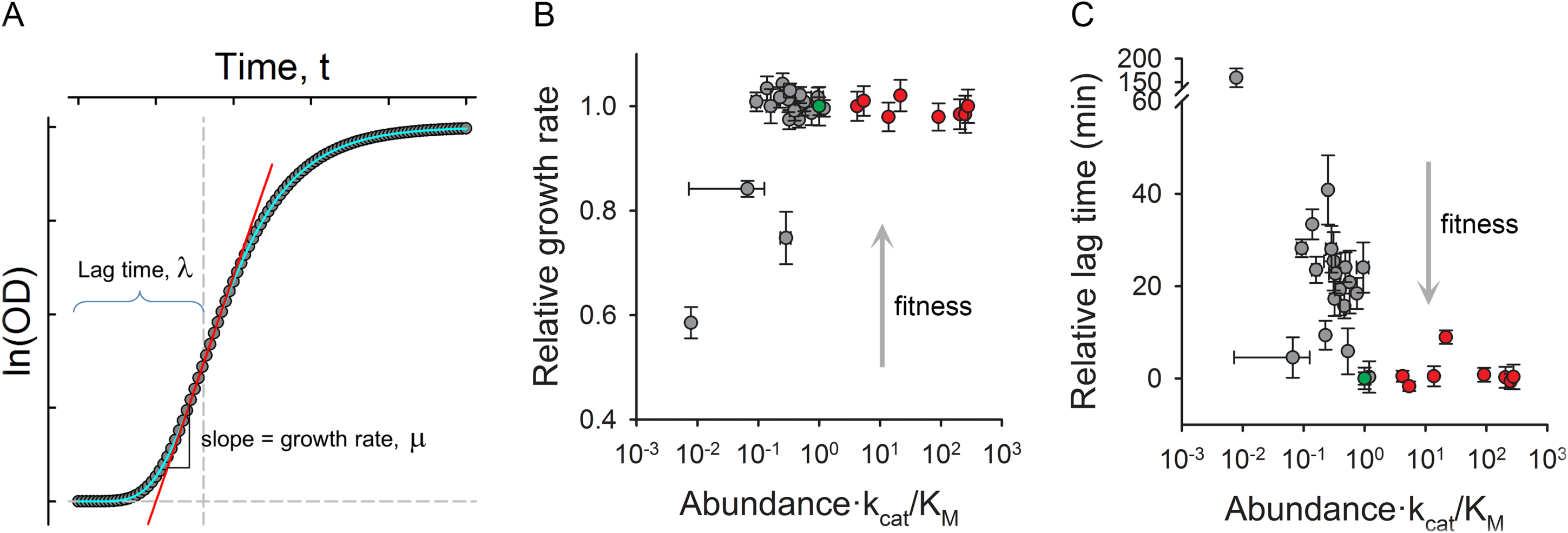
Traits of population growth. (A) Schematic of estimation of lag time and growth rate. The representative data points (solid gray circles) were plotted as ln(OD) vs time and was fitted to a four parameter Gompertz function (Eq. 2) (cyan line). The red line is a tangent at the inflection point of the function. The slope of the tangent is considered as the growth rate ^(*μ*)^ and the time required to reach the maximum growth rate or the inflection point is taken as the lag time ^(*λ*)^ (vertical dashed line). (B) Relative growth rate (μ/μ_*WT*_) and (C) relative lag time (λ−λ_*WT*_) as functions of catalytic capacity which is defined as abundance ×*k*_*cat*_/*K*_*M*_. The mutant data is shown in gray circles, whereas red circles represent the BW27783 strain with varying degrees of overexpression of WT Adk from a pBAD plasmid. Data for WT is shown in green. The data points represent mean and error bars represent standard deviation of parameters derived from growth curves of 3 colonies (biological replicates) in triplicates (9 curves). See related Figs. S6-S11 and Tables S2-S3. The solid gray arrow indicates the direction of increasing fitness.

### WT E. coli is positioned at the cusp of the biophysical fitness landscape for lag time

Since almost all the designed mutants were destabilizing and therefore have lower catalytic capacity than *E. coli* WT, they only provide sampling in the lower range of catalytic capacity. In studies so far, no evidence exists for changes in intracellular protein abundance for stabilizing mutations. Hence to determine the dependence of growth rate and lag time on catalytic capacity above WT levels, we over-expressed WT Adk from a pBAD plasmid (see Supplementary Methods). We observed no significant change in either growth rate or lag time at higher than endogenous catalytic capacity (Fig. 2B, C, S8, and Table S3). This means that while the growth rate appears to be insensitive to large changes in Adk catalytic capacity both below and above the wild-type level, WT catalytic capacity appears to be situated at the threshold of optimizing lag time. Next, we attempted to quantitatively compare the position of WT on these two fitness landscapes. To that end, we used a simple reciprocal Michaelis-Menten-like function to fit the relative growth times (growth time is reciprocal of growth rate ^*μ*^) and lag times (Fig. S11, also see Eq. 3 and *Methods*). The fitting parameter ^*b*^ which characterizes the onset of curvature on the landscape (analogous to ^*K*_*M*_^ in Michaelis-Menten equation for enzymatic rate) reports proximity of WT to the cusp on the landscape (see *Methods*). It was 0.006 for growth time and 0.019 for lag time as compared to normalized catalytic capacity of 1 for WT. This shows that WT is situated closer to the cusp in terms of lag time as compared to growth time or growth rate.

### Shorter lag imparts advantage at low carrying capacity: A computational model

This data highlights the pleiotropic effects of mutations on different phases of bacterial population growth, which raises the question of how pleiotropy shapes the evolutionary fate of a mutation. We explore this issue by considering the outcome of binary competitions between strains^18^. We first simulated binary competitions over a wide range of growth rates and lag times in media conditions that allow for either 5-fold (low carrying capacity) or 500-fold (high carrying capacity) increase over the initial population (Fig. 3A) (See *Methods*). We found that there is a significant tradeoff between lag times and growth rates in determining the winners of binary competitions, with lag playing a more important role at low carrying capacity (Fig. 3A), implying that beneficial lag provides a greater fitness advantage under strongly nutrient-limiting conditions.

**Fig. 3:**
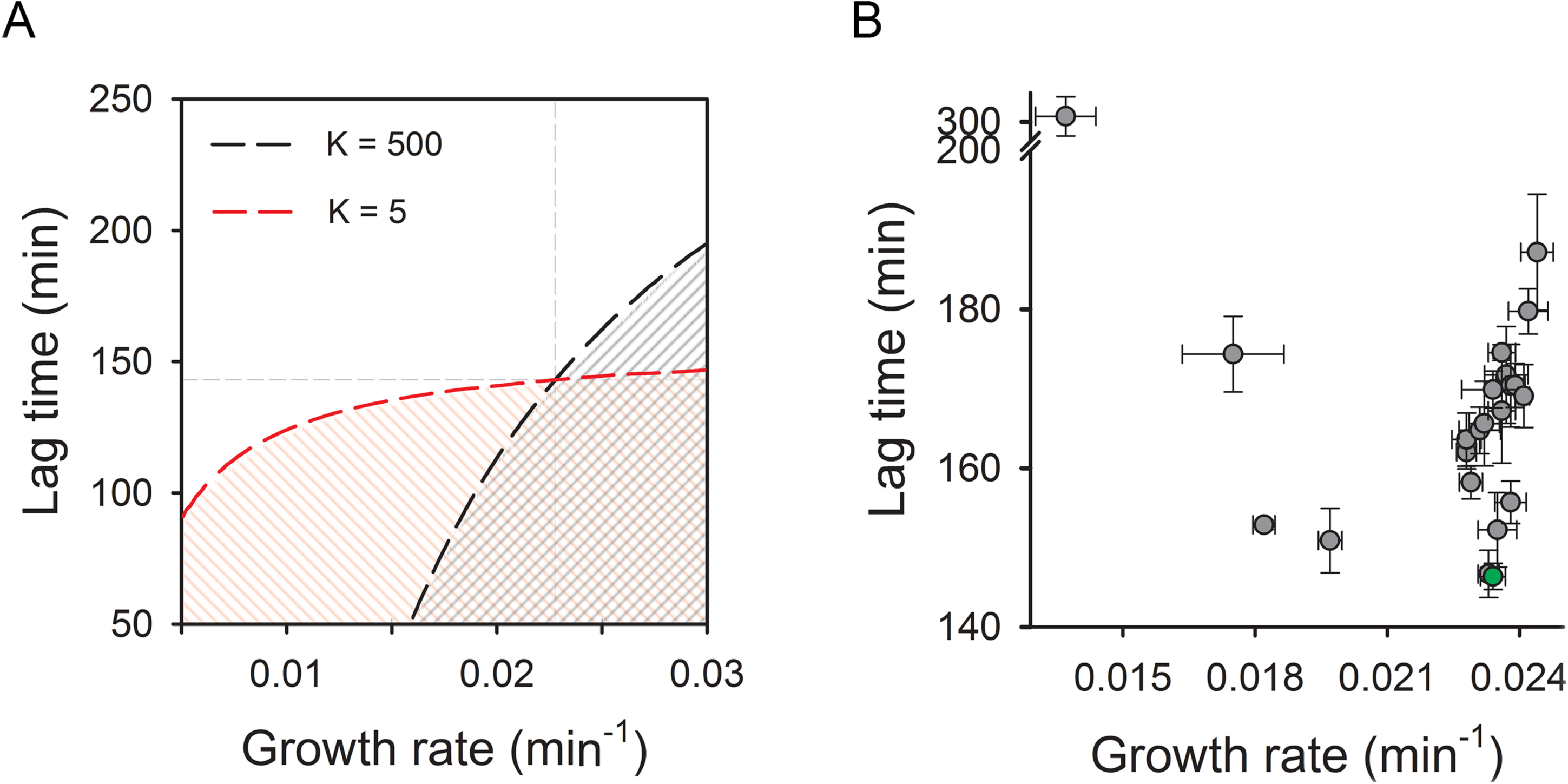
Binary growth competition. The growth of individual strains was modeled as per Gompertz equation (Eq. 2). The growth parameters for strain 1 were fixed to those obtained for WT Adk (dashed gray lines) while those for strain 2 were generated randomly over a wide range of growth rates (0.005 to 0.030 min^-1^) and lag times (50 to 250 min). (A) Contour plot showing fraction of strain 1 (WT) at saturation when the competition is carried out under two different carrying capacities (red line indicates *K* = 5 while the black line indicates *K* = 500). The dashed lines indicate neutrality region where both strains have equal proportions at saturation. The areas below the neutrality line (filled with solid lines) represent the parameter space where strain 2 wins the competition (fraction of strain 2 > 0.5). (B) Scatter plot of growth rate ^(*μ*)^ versus lag time ^(*λ*)^. The data points represent the mean and error bars the standard deviation of 6 to 9 measurements (see Table S2). The growth rate and lag time appear to be statistically independent of each other across the Adk mutant strains (Spearman’s *ρ* = 0.31, *p* = 0.15).

### Shorter lag imparts advantage at low carrying capacity: Experimental evidence

To realize varying nutrient conditions in binary competition experiments, we explored the growth of *E. coli* over a range of glucose concentrations, mimicking the variation of carrying capacity in simulations, and found that only the carrying capacities are proportional to glucose concentration with minimal effects on lag time and growth rate (Fig. 4). This suggests that observing the outcome of the competition at different time snapshots in a nutrient-rich medium is equivalent to running the competition at different glucose concentrations (carrying capacities). To evaluate the predictions from simulations, we carried out two sets of binary competition experiments based on the overall distribution of growth rates and lag times (Fig. 3B). First, we selected strains exhibiting a tradeoff between growth rate (μ) and lag time (λ) (μ_1_ >μ_2_ and λ_,1_>λ_,2_ (inset of Fig. 5B). Second, we tested competition between strains that differ in their lag times but have nearly indistinguishable growth rates (μ_1_ ≈ μ_2_ and λ_1_ >λ_2_ (inset of Fig. 5C). In all cases a strain with shorter lag time is expected to dominate at lower carrying capacity conditions (corresponding to the competition outcome at early time points), however this advantage would be lost at later time points if its growth rate is lower than that of the competing strain (Fig. 5A). In the second scenario, the advantage due to short lag is expected to persist even at high carrying capacity conditions because the growth rates of the competing strains do not differ. We estimated the relative proportions of the two strains by a qPCR-based mismatch amplification mutation assay (MAMA) approach^19^ (see *Methods* and Fig. S12). As expected in the first scenario, L083F and V106H dominated at earlier time points when competed against A093I and L209I, respectively, due to their shorter lag times (λ_L083F_ < λ_A093I_ and λ_V106H_ < λ_L209I_) (Fig. 5B). Eventually their fraction dropped below 0.5 at later time points (equivalent to high carrying capacity) where the growth rates determine the competition output (μ_L083F_ < μ_A093I_ and μ_V106H_ < μ_L209I_) (Fig. 5B). Similarly, for the second scenario, despite having similar growth rates (μ_*WT*_ ≈ μ_Y182V_ ≈ μ_L209A_), the fraction of WT was always maintained above 0.5 as it spends a shorter time in the lag phase compared to Y182V and L209A (Fig. 5C). The early advantage to WT due to its shorter lag phase determined the competition fitness throughout the whole growth cycle.

**Fig. 4:**
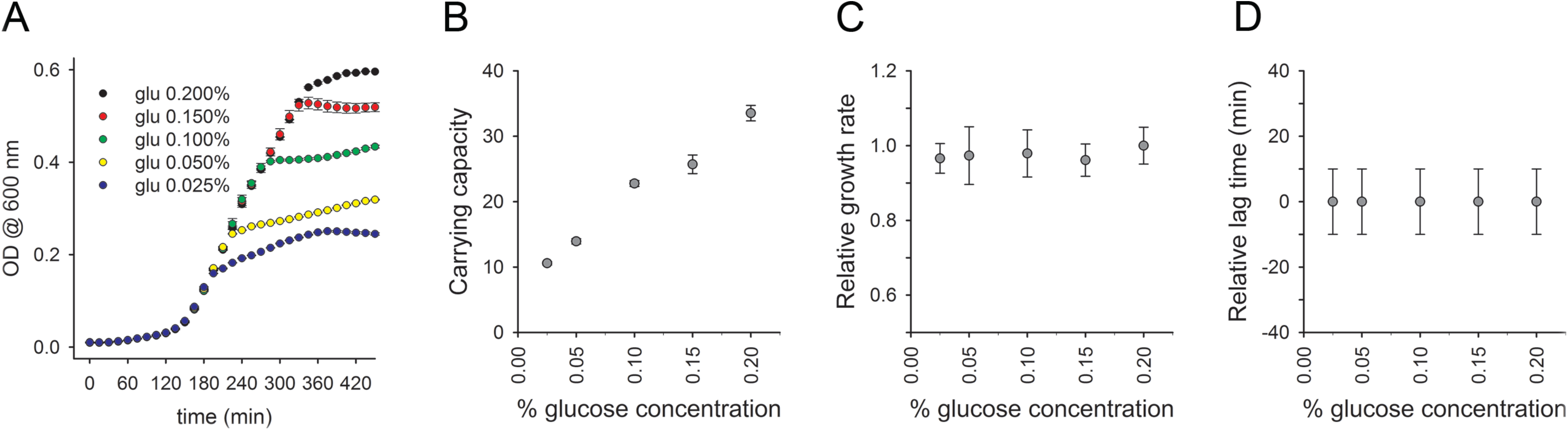
Growth curves at various nutrient concentration. (A) Growth curves of strains with WT Adk obtained under varying glucose concentrations in supplemented M9 medium. The fitted growth curve parameters are shown as functions of glucose concentration: (B) carrying capacity of ln(OD) as derived from Gompertz fitting, (C) relative growth rate (μ/μ_0.2_), and (D) relative lag time (λ-λ_0.2_). The growth rates and lag times are estimated from analysis of growth curve derivatives and are normalized relative to the respective values at 0.2% glucose concentration.

**Fig. 5:**
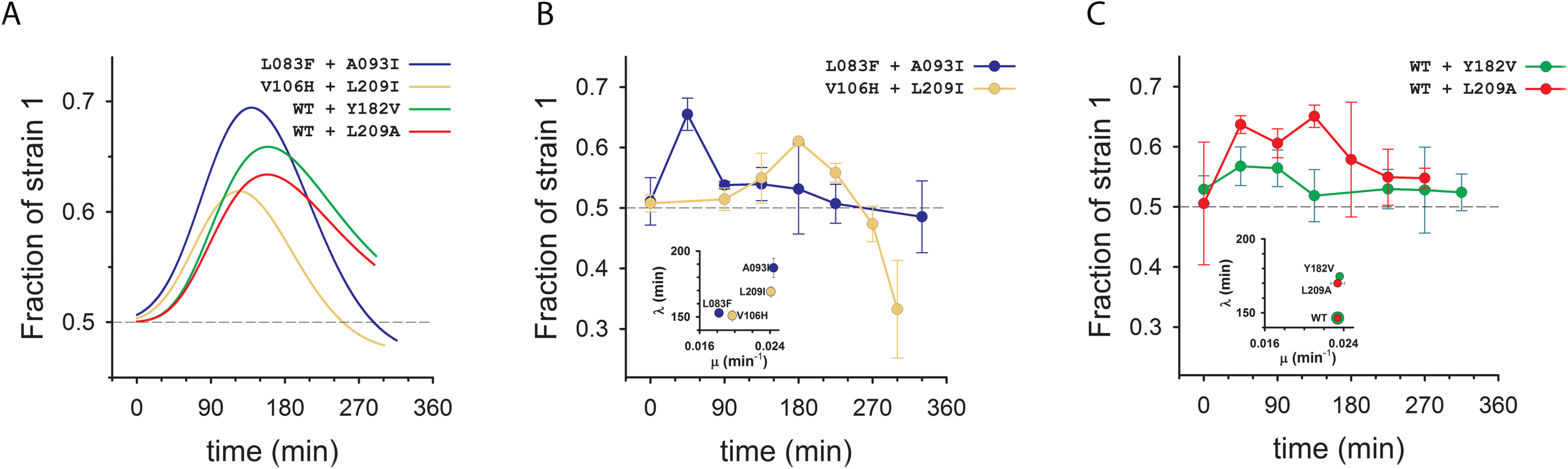
Tradeoffs between lag and exponential growth in binary competitions. (A) Fraction of the first strain as a function of time in simulated binary competitions. We modeled growth of each strain using the Gompertz 4-parameter equation (Eq. 2) with experimentally measured growth rate and lag time values. The initial OD for individual strains was assumed to be 0.006 at the start of competition, and growth was assumed to saturate at OD of 0.6. Despite having similar growth rates, the fraction of WT in WT + L209I and WT + Y182V competitions was always above 0.5 owing to the advantage it gained due to shorter lag time (scenario 2 in the text). L083F and V106H dominate at earlier time points (equivalent to low carrying capacities) due to their short lag times compared to their respective competitors. However, at longer times (high carrying capacities) the advantage due to lag is lost due to their lower growth rates. (B, C) Experimental validations of the predictions in (A) using qPCR based mismatch amplification mutation assay (MAMA). The fraction of competing strains was estimated using Eq. 4. The data points are mean and error bars represent standard deviation of two measurements. See related Fig. S12. The growth rates and lag times for the competing pairs are shown in insets.

## Discussion

A complete mapping of mutational fitness effects would require sampling a practically infinite number of mutations, an impossible proposition. Instead, we can project fitness onto a fairly small number of molecular properties of proteins^5–7,20^. Within this paradigm, the identity of a particular mutation does not matter as much as its effect on essential biochemical and biophysical properties of the proteins in question. Our 21 engineered mutations in Adk, along with the overexpression data, allow us to outline the biophysical fitness landscape, covering a wide range of variation of the physical parameters of the Adk protein. This data shows that we can collapse several molecular phenotypes into a single effective parameter – the product of protein abundance and activity *k*_*cat*_/*K*_*M*_ (catalytic capacity) – which quantitatively determines the biophysical fitness landscape to a great extent (Fig. 2B, C). That is, Fig. 2 indicates that the fitness effects of mutations can largely be predicted from their biophysical effects over a broad range of catalytic capacity. Indeed, Adk catalytic capacity explains the variation in lag times to a large extent (Fig. S10), validating the utility of a biophysical fitness landscape for mapping fitness effects.

These results illustrate how the evolutionary endpoint of molecular traits may depend fundamentally on the multidimensional nature of fitness, with the relative importance of different components of fitness depending on the environment and lifestyle of the organism. It has been argued that endogenous molecular traits are established as a result of mutation-selection balance^21^, with the final outcome depending on the relative strengths of selection and genetic drift as determined by the population structure^22,23^. Here we encounter a more complex situation where mutations in the essential enzyme Adk change multiple fitness components. In this case, the mutation-selection balance apparently resulted in disparate outcomes for the two fitness components with respect to the molecular trait, placing lag time at the cusp while keeping the exponential growth rate farther within the plateau region of its respective biophysical fitness landscape. Such an outcome may reflect different strengths of selection on growth and lag. The relative strength of selection on these fitness components depends crucially on the environmental conditions (e.g. nutrient availability, etc.)^24^. Our studies of binary competitions (Figs. 3 and 5) highlight this scenario by showing how the environmental parameter of carrying capacity can determine winners and losers in evolutionary dynamics. Although the lag time of a population can depend not only on the environment but also on the population’s specific history (e.g., how long it was previously in stationary phase), the fundamental role of Adk in metabolism suggests that its effects on lag time are likely to be common across conditions and histories. The deep connection between ecological history of species and optimization of biophysical traits of their proteins is a subject for valuable future studies.

Much of our current understanding of microbial cultures and fitness comes from experiments done in the laboratory, where strains are typically grown under a large supply of nutrients. The situation might be very different in the wild, however, where bacteria and other microbes have to survive under harsh conditions of nutrient starvation, extreme temperature, and other environmental stresses^25–27^. For example, *E. coli* is the predominant facultative anaerobe in the gastrointestinal tract^28^ which allows it to thrive in fluctuating environments of differing oxygen concentrations along the GI tract (e.g., the small vs. the large intestine). In these circumstances, organisms are likely to spend only a minute fraction of their life-cycle in the exponential growth phase, while undergoing many cycles of lag-growth-saturation as new resources become available and old ones are exhausted. It is therefore intuitive to expect that there has been strong selection in favor of organisms that can not only divide rapidly during exponential growth, but that can also wake up quickly from their lag phase and respond to newly available resources. Our study demonstrates how this selection may shape individual molecular traits.

This work highlights the relationship between various components of fitness and the molecular properties of modern enzymes — the endpoint of evolutionary selection. An interesting question which is beyond the scope of current work is how modern variants emerged in evolutionary dynamics. To that end mapping temporary reconstructed ancestral species onto biophysical fitness landscape of Adk (and other enzymes) appears a promising direction of future research.

## Methods

Selection of mutations: Mutations at relatively-buried positions generally result in decreased stability and lower fitness^13,29^. Hence we selected the sites for mutagenesis with side-chain accessibility of less than 10%. In addition, the selected sites were also away from the active-site residues, or active-site contacting residues, and a minimum of 6 Å away from the inhibitor Ap5A binding sites (pdb 1ake). The structure of Adk is divided into three domains: LID (residues 118-160), NMP (residues 30-67), and Core (residues 1-29, 68-117, and 161-214). We define the active-site residues as those whose accessible surface area changes by at least 5 Å^2^ in the presence of the inhibitor Ap5A. A similar criterion was used to define the residues contacting the active site. Altogether 4 residues from the LID domain, 3 from the NMP domain, and 28 from the Core domain satisfy these criteria. Of the 28 sites from the Core domain, we randomly chose 6 to mutate. We chose the identities of the mutations to span various sizes of the side chains and a range of conservation. We derived the conservation from the multiple sequence alignment of 895 sequences for Adk collated from ExPASy database (as of Nov 2012).

Generation of mutant strains: We generated the strains with WT and mutant *adk* with chloramphenicol- and kanamycin-resistance genes on either end of the *adk* gene using the genome editing approach as described previously^3^. Since the *adk* gene is flanked by two repeat regions (REPt44 and REPt45) on the wild-type chromosome, we extended the homology required for recombination up to the middle of the adjacent genes.

Growth curve measurements and media conditions: WT and mutant strains were grown overnight at 30 °C from single colonies in a supplemented M9 medium (0.2 % glucose, 1 mM MgSO4, 0.1 % casamino acids, and 0.5 μg/ml thiamine). OD600 was measured for all the strains and then the cultures were normalized to whichever had the lowest OD. The normalized cultures were diluted 1:100 in fresh supplemented M9 media and the growth curves were monitored in triplicates using Bioscreen C at 37 °C. We derived the growth parameters by fitting ln(OD) versus time with the four-parameter Gompertz function (see below). The error in replicates was found to be between 2-3% on an average, and it did not improve significantly upon increase in number of replicates.

Fitting growth data and estimation of growth parameters: In our study, we define lag time ^(*λ*)^ as the time required to achieve the maximum growth rate ^(*μ*)^ (Fig. 2A). Growth time ^(*τ*)^ was defined as reciprocal of growth rate ^*μ*^. Since it has the same units as lag time, it is more convenient to use for the statistical analysis and data fitting (Fig. S11).

We used two different methods to infer these parameters: A) direct analysis of growth curve derivatives and B) fits to the Gompertz function (Fig. S6).

In method A, we took the growth rate as the maximum value of 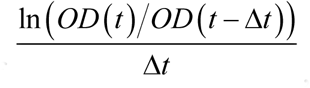 where Δ*t* is 15 minutes. The lag time was then the earliest time at which this maximum growth rate was achieved.

For method B we used the following four-parameter Gompertz function to fit ln(OD) vs. time plots:

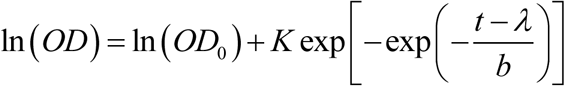

where the carrying capacity is ^*K*^, the maximum growth rate is μ=*K*/(*b*·exp(1)) and the lag time ^*λ*^ is the time taken to achieve the maximum growth rate.

For both the methods, we considered only data points with *OD*600 ≥ 0.02. The instantaneous derivatives of all growth curves show presence of a distinct peak at OD600 values greater than 0.02 (Fig. S6), indicating monoauxic growth and also asserting that the derived growth parameters are unaffected due to ignoring the lower OD data.

The ^*μ*^ and ^*λ*^ estimated from the two aforementioned methods are strongly correlated (Pearson’s *r* = 0.80, *p* = 1.4e−5 for ^*μ*^ and *r* = 0.71, *p* = 3.0e-4 for ^*λ*^) (Fig. S7). However, the uncertainty in the fitted parameters appears to be less than the uncertainty in the parameters obtained from the derivatives, which are limited by the low time-resolution of the experimental data (acquired at an interval of 15 min).

The growth rate ^(*μ*)^ and lag time ^(*λ*)^ appear to be statistically independent of each other across the Adk mutant strains (Spearman’s *ρ* = 0.31, *p* = 0.15, Fig. 3B). Hence it is conceivable that selection can act separately on these two traits, which is further illustrated by the different fitness landscapes observed when projected onto the axis of catalytic capacity (Fig. 2B, C).

Statistical tests for mutational variation in growth and lag phases: We estimated the monotonic relationship between various growth traits and molecular/cellular properties of Adk mutant proteins using Spearman’s rank correlation (Fig. S10). The agreement between growth parameters derived using instantaneous derivatives and Gompertz fit were estimated by Pearson’s correlation coefficient (Fig. S7). We excluded V106N from all statistical analysis and data fitting as its lag time is ~13 s.d. away from the average lag time of all other strains.

Quantification of the location of WT on the fitness landscapes: A Michaelis-Menten-like elasticity curve function has been used previously^5,6,16,20^ to fit the dependence of growth rate on catalytic capacity. Since we are considering growth and lag times rather than rates, we use a reciprocal form of the Michaelis-Menten-like function for fitting relative growth time (τ/τ_*WT*_) and relative lag time (λ/λ_*WT*_) vs. catalytic capacity (Fig. S11):

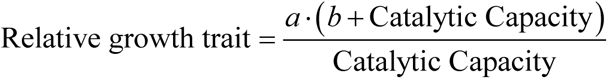

where ^*a*^ is the asymptotic value of the trait for infinitely large catalytic capacity, and ^*b*^ is the catalytic capacity when the trait equals twice the asymptotic value ^(2*a*)^. Since catalytic capacity is normalized by WT, ^*b*^ serves as a measure of how close to the cusp the WT on the respective landscapes is. For fits in Fig. S11, we empirically set ^*a* = 1^ which enables easy comparison of parameter ^*b*^ for lag time and growth time plots.

Simulation of binary competition: We simulated the competition of two strains by using the Gompertz function (Eq. 2) to model the growth of individual strains. The initial population (OD0) for both strains was equal, and growth ceases when ∑(*OD*_*t*_ *OD*_0_)_*i*_=*K* where *K* is the carrying capacity. We considered two different values of carrying capacities (5 and 500). We set ^*μ*_1_^ and ^*λ*_1_^ to values derived experimentally for WT Adk strain (Table S2), while the growth rates and lag times for the second competing strain were varied randomly across the intervals 0.005 to 0.030 min^-1^ (for growth rate) and 50 to 300 min (for lag time).

Binary growth competition and quantification: The overnight cultures for individual strains were grown for 16 hours at 30 °C. These cultures were mixed in 1:1 proportion, diluted to an OD of 0.01 in fresh supplemented M9 media, and then regrown at 37 °C. The samples were drawn at different time points, and the OD was adjusted to 2.0, either by concentration or dilution. 5 µl of OD 2.0 culture was eventually diluted in 45 µl of lysis solution (QuickExtract DNA extraction solution (Epicentre)) to reach OD 0.2. Genomic DNA extracted from 50 µl of OD 0.2 culture was diluted 5000 times and used as template. The individual strains in the competition were differentially amplified using allele-specific primers and quantified by a qPCR-based mismatch amplification mutation assay method ^19^ using QuantiTect SYBR Green PCR kit (Qiagen). A 150 bp long non-mutagenic amplicon of *adk* gene was amplified as a reference to quantify total genomic DNA. The fraction of the competing strains was determined using the following equation:

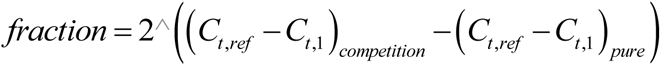

where ^*C_t_*^ represents threshold cycle of qPCR, ^*ref*^ and 1 are the PCR reactions for amplifying the reference and the first allele in competition, while ^*competition*^ and ^*pure*^ represent the condition of culture.

Data availability: All raw data for growth curves of *adk* WT and mutant strains, as well as WT overexpression in *E. coli* BW27783 strains, is included as Dataset 1.

## Acknowledgements

We thank Shimon Bershtein and Adrian Serohijos for helpful discussions.

### Contributions

BVA and EIS - designed research; BVA, SB, JT and MMu - performed experiments; BVA, MMa, SB and EIS - analyzed the data; BVA, MMa, SB and EIS - wrote the paper; All authors edited and approved the final version.

## Supplementary Methods

### Mutagenesis and protein purification

Adenylate kinase (Adk) is encoded by the *adk* gene, which was cloned under the T7-*lac* promoter in pET28a(+) vector (Invitrogen) between *Nde*I and *Xho*I restriction sites. We carried out mutagenesis with a pair of 30-35 bp long, partially-complementary primers and the inverse PCR technique using KOD hot-start DNA polymerase. The mutations were centered in the complementary regions of the primers. The mutagenic plasmids were transformed in *E. coli* DH5α cells for faithful propagation and storage, and in *E. coli* BL21(DE3) for protein overexpression and purification. The His-tagged proteins were purified by Ni-NTA affinity chromatography (Qiagen) and subsequently passed through a HiLoad Superdex 75 pg column (GE). The monomeric peak was collected, concentrated and eventually stored in 10 mM potassium phosphate buffer (pH 7.2). The concentration of the proteins was measured by BCA assay (ThermoScientific) with BSA as standard.

### Biophysical characterization

Thermal denaturation: We assessed the thermal stability of WT and mutant proteins by differential scanning calorimetry (nanoDSC, TA instruments) using 20 μM of protein. The scans were carried out from 10 to 90 °C at a scan rate of 90 °C/hr. The thermodynamic parameters were derived by fitting the data to a two-state unfolding model using NanoAnalyze (TA instruments). We also carried out thermal denaturation using the melt-curve module of BioRad CFX96, with Sypro Orange dye as a probe for unfolding as described earlier^1^. The dye was added to the final concentration of 5× in a 25 μl reaction volume containing 4 μM of protein in 10 mM potassium phosphate buffer (pH 7.2). The data were fit to a standard four-parameter sigmoidal equation to obtain apparent melting temperatures.

Urea denaturation: We carried out isothermal urea denaturation with WT and mutant proteins to assess the stability of the proteins to chemical denaturants. We incubated 5 μM of protein for ~4 hrs at 25 °C with varying concentrations of urea (0-8 M). The urea concentrations were estimated by refractive index measurements. The denaturation was monitored by measuring the ellipticity at 222 nm using a CD spectrometer (Jasco). The melt data was fitted assuming a model of two-state unfolding with linear free energy as described earlier^2,3^. The m-value was fixed to 3300 cal/mol/M for fitting.

Gel filtration: We assessed the oligomeric status of purified proteins by gel filtration using 50 μg of protein on sephadex 75 analytical columns.

ANS and proteostat binding: We used 12 μM of bisANS for assessing binding to 2 μM of protein in 10 mM potassium phosphate buffer (pH 7.2). The excitation and emission wavelengths were set to 395 nm and 490 nm, respectively. 2 μM of protein was incubated with 3.5 mM of the proteostat dye in 1× assay buffer (Enzo LifeSciences). For this the excitation and emission wavelengths were set to 550 and 600 nm, respectively.

Enzyme activity: We measured the activity of Adk in terms of ADP formation by an end-point assay as described earlier^4^. Briefly, the concentration of AMP was fixed to 500 μM and ATP concentration was varied from 0 to 500 μM in an enzymatic reaction. 5 nM of Adk was used to initiate the reaction and 500 μM of Ap5A was used for quenching at 20, 40, and 60 second time points. The amount of ADP formed was measured by LDH-Pyruvate kinase-coupled reaction and the kinetic parameters were derived by fitting the data to the Michaelis-Menten equation.

Adk overexpression: The *adk* gene was cloned in a pBAD plasmid and transformed in the *E. coli* BW27783 strain (CGSC#12119). This strain constitutively expresses the arabinose transporter (araE) which enables uniform uptake of arabinose. The cells were induced with increasing concentrations of arabinose from 0 to 0.05%.

Intracellular protein abundance: Cells were grown in supplemented M9 medium for 4 hours at 37 °C, harvested and subsequently lysed with 1× BugBuster (Novagen) and 25 units/ml of Benzonase. Total amount of proteins in cell lysate was estimated by BCA assay. The specific fraction of Adk was determined by SDS-PAGE followed by western blot using rabbit anti-Adk polyclonal antibodies (custom-raised by Pacific Immunology).

Estimation of viable cells in saturating culture: The overnight culture was grown in supplemented M9 medium for 16 hours at 30 °C and the proportion of live:dead cells was measured using Live/Dead BacLight Bacterial Viability Kits (Molecular Probes) according to the manufacturer’s instructions. Briefly, 1×10^8^ cells (in a volume of 1ml) were mixed with 3 µl of a 1:1 proportion of Syto9 dye and Propidium Iodide (PI). The mixture was incubated in the dark for 15 minutes, following which the fluorescence was measured at 530 nm and 630 nm. Syto9 dye stains live cells and emits fluorescence at 530 nm (green), while PI stains dead cells and can be detected at 630 nm (red). The ratio of fluorescence values at 530 nm:630 nm corresponds to the proportion of live:dead cells in that sample which was eventually used to estimate the percentage of live cells in a sample, according to the manufacturer’s instructions. An exponentially growing culture (considered as 100% live) and cells treated with 70% ethanol for 1 hour (considered 100% dead) were mixed in different known proportions, and their 530:630 nm ratio was used to generate a standard curve.

**Fig S1:**
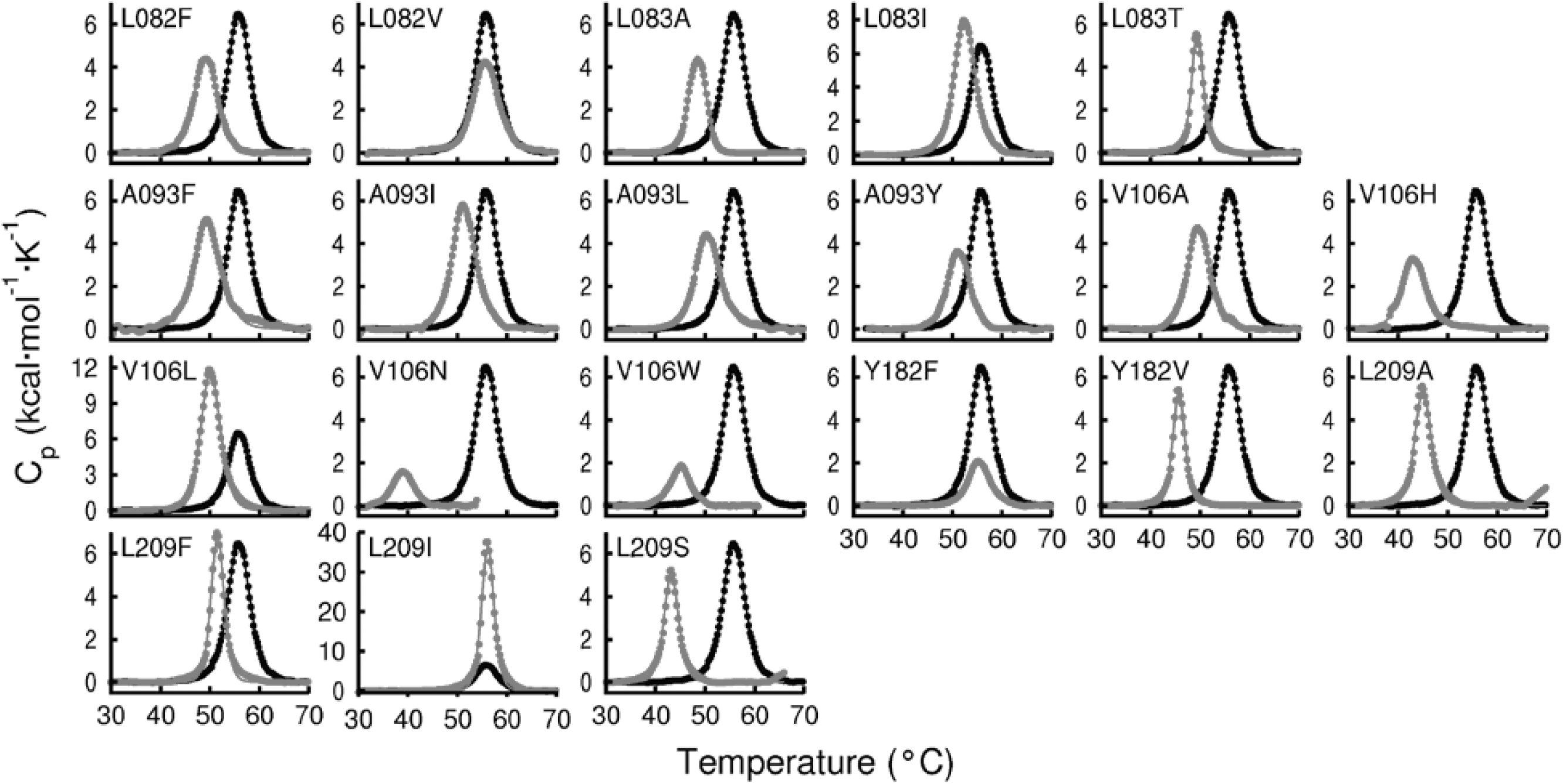
Thermal unfolding monitored by Differential Scanning Calorimetry (DSC) for WT (black trace) and 20 different Adk mutant proteins (red trace). The molar heat capacity (C_p_) is shown as a function of temperature. The scan rate was 90 °C/hr. The data was fitted to a two-state thermal unfolding model to derive the thermodynamic parameters.

**Fig S2:**
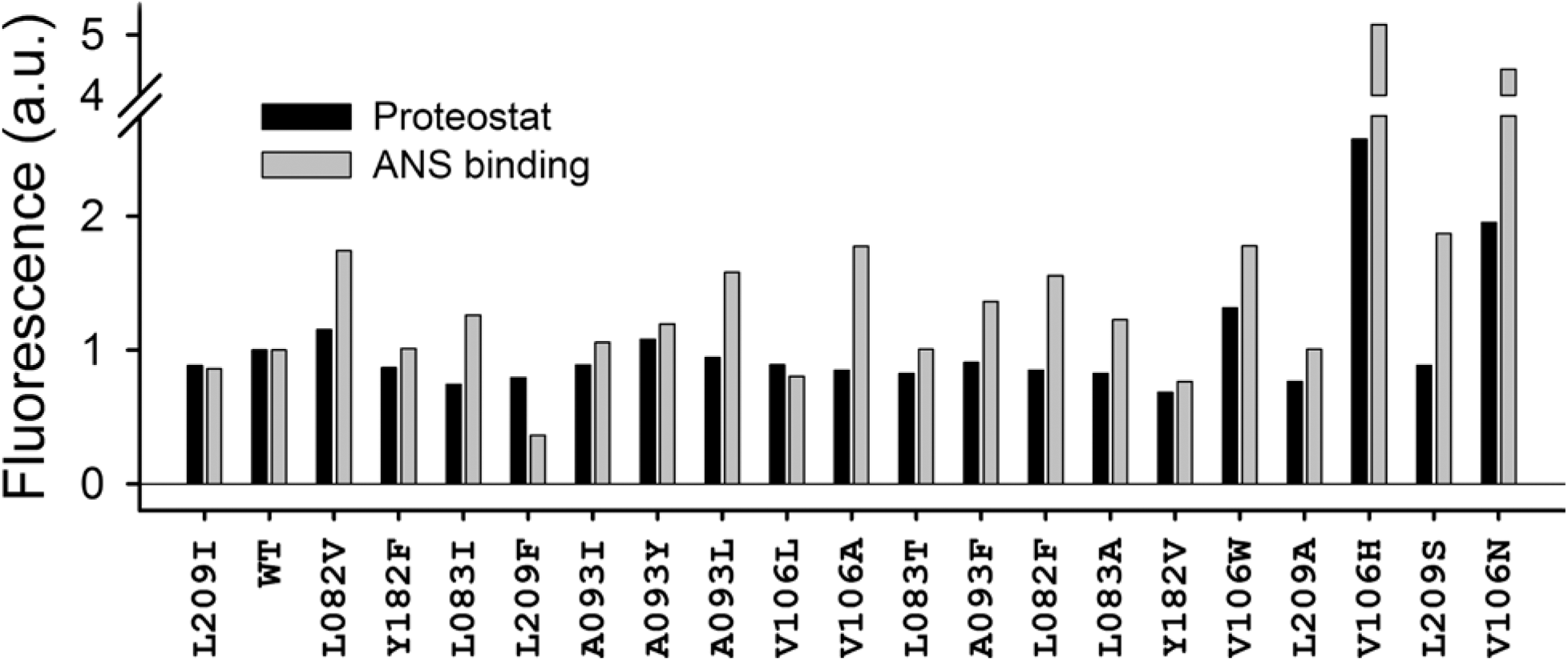
Isothermal urea denaturation curves at 25 °C for WT (black dots) and mutant Adk proteins (red dots). The fraction unfolded (Fu) is plotted as a function of denaturant concentration. Protein denaturation was monitored by recording the CD signal at 222 nm. The data was fit to a two-state unfolding model.

**Fig S3:**
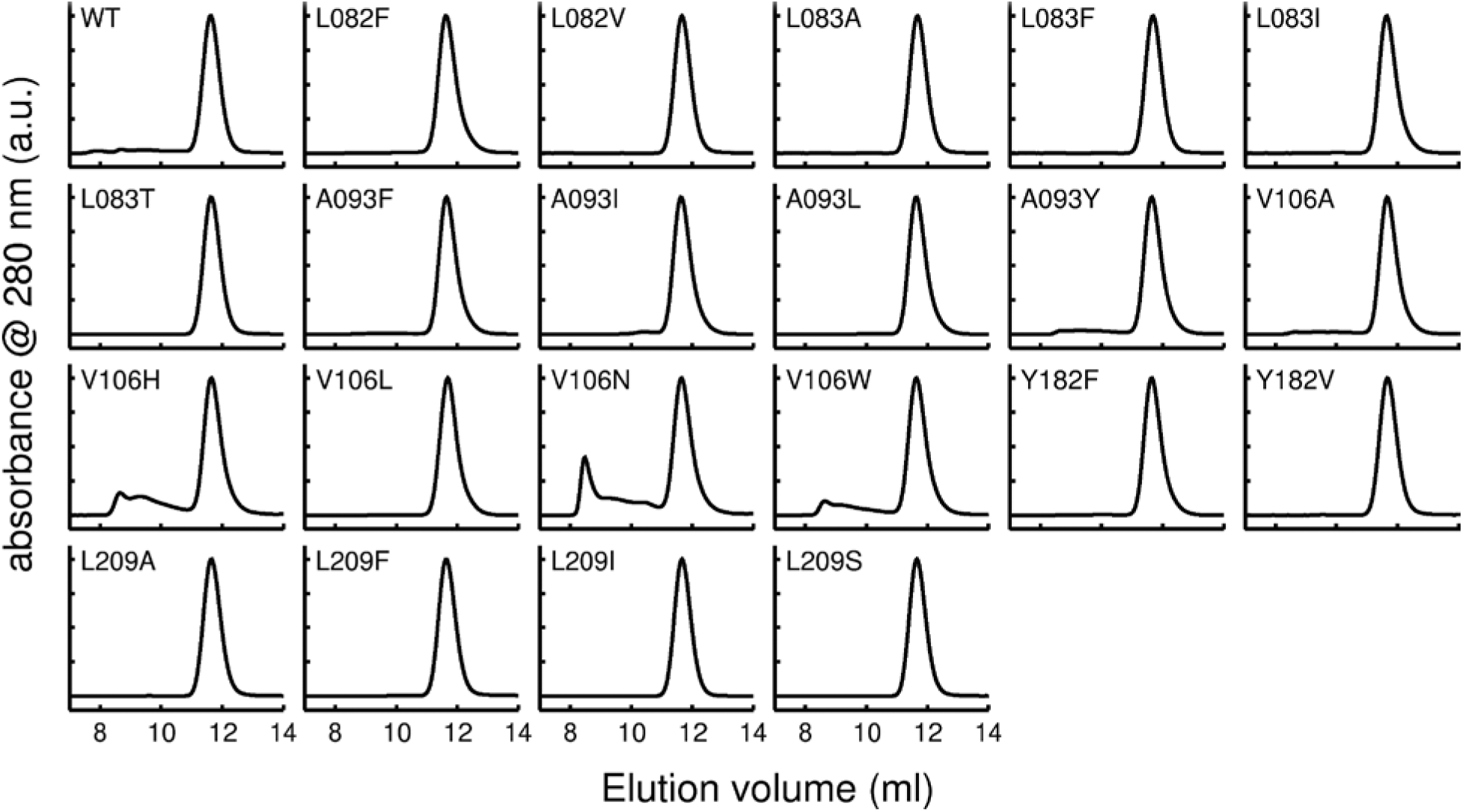
Analytical gel-filtration profile of WT and 20 mutant Adk proteins on a Superdex-75 column at room-temperature. The absorbance at 280 nm is shown as a function of elution volume. For comparison all the monomeric peaks were normalized to 1. WT Adk along with most other mutant proteins elutes at the expected position for a monomer. Exceptions were V106H, V106N and V106W, where additional peaks appear at much higher molecular weights.

**Fig S4:**
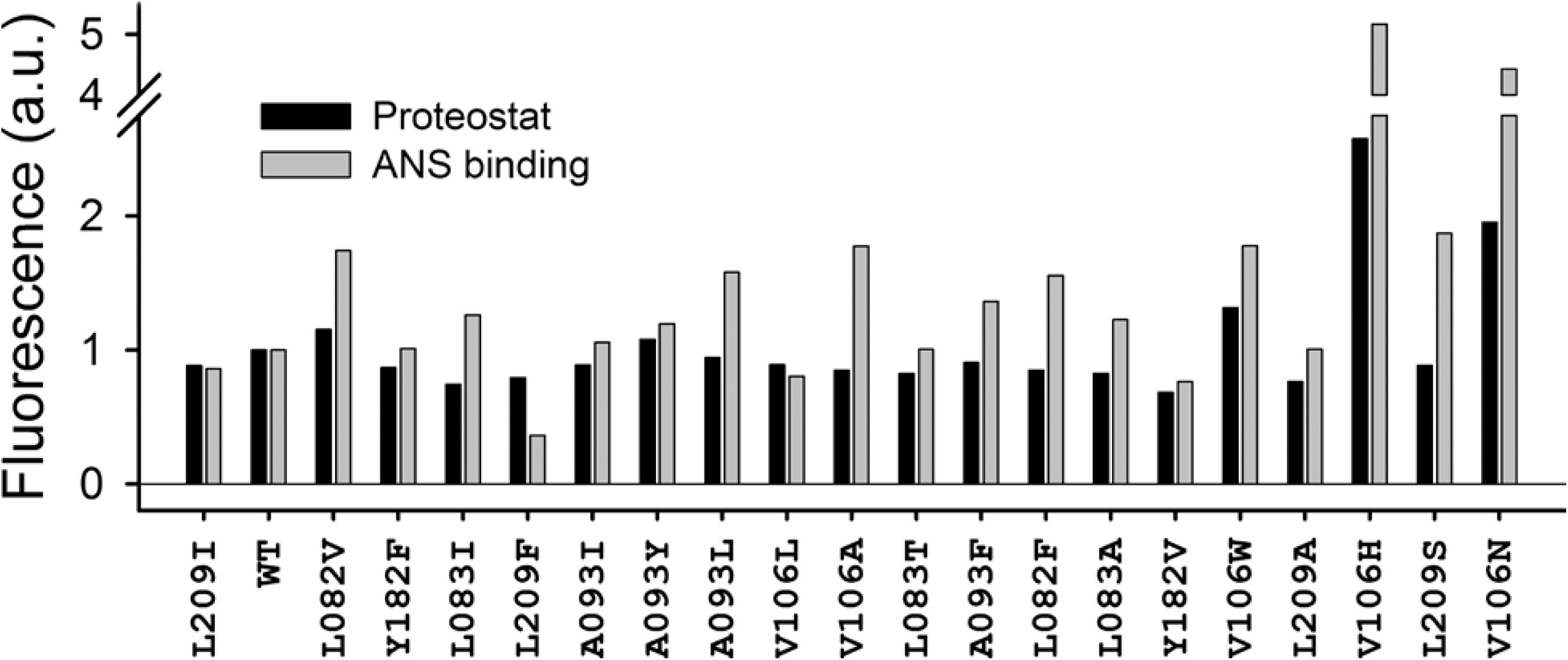
Aggregation propensity and molten-globule states of mutant proteins. Bar plots represent the extent of ProteoStat and ANS binding to WT and mutant Adk proteins. The proteins on the x-axis are arranged in decreasing order of stability from left to right.

**Fig S5:**
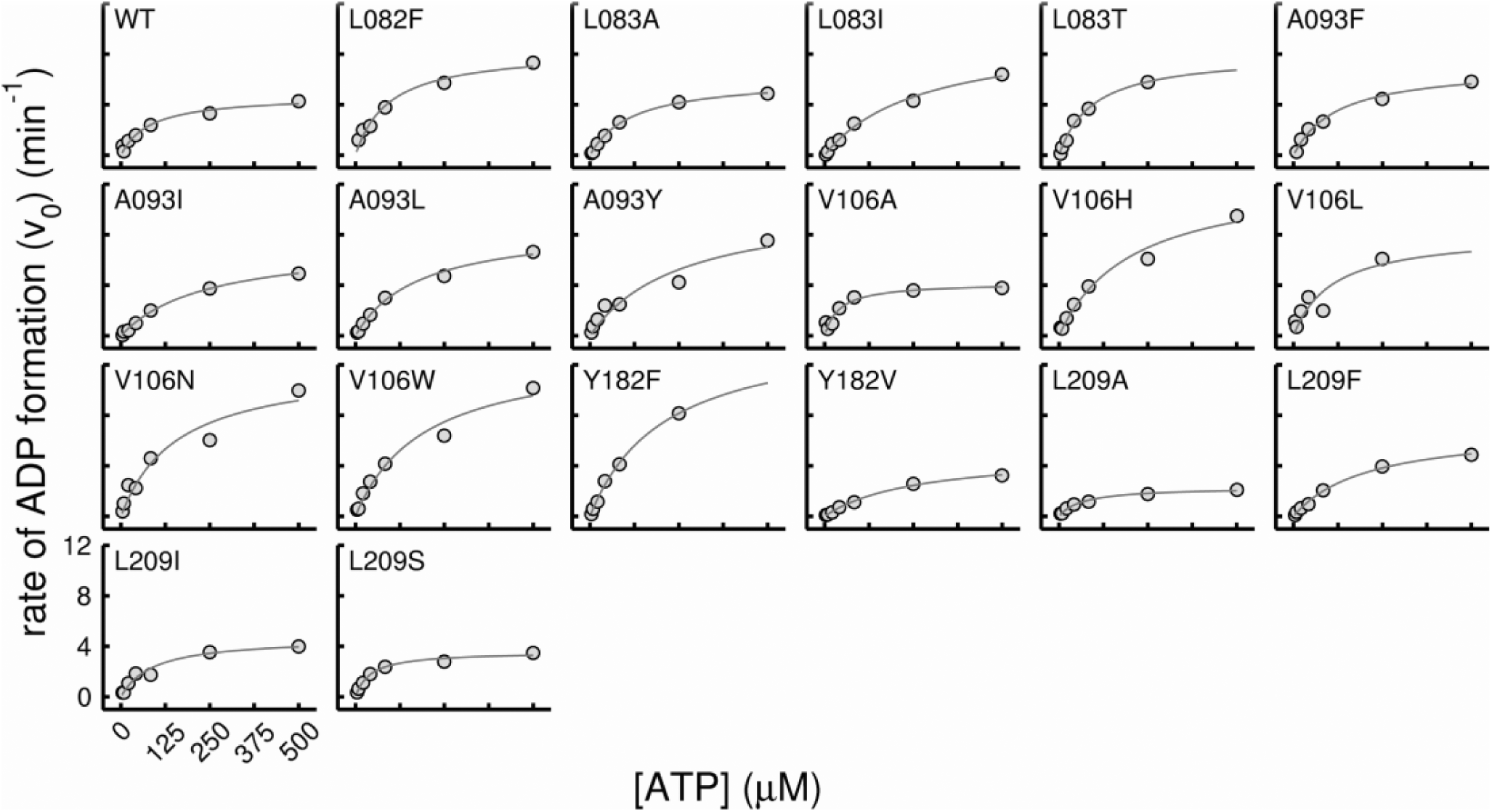
Enzyme activity of Adk mutants at 25 °C measured as described in Supplementary methods. The initial velocity, shown as a function of ATP concentration, was calculated as the amount of ADP produced per minute by 1 nmol of Adenylate Kinase. The concentration of AMP in all experiments was fixed to 500 µM. The data (gray circles) was fitted using the Michaelis-Menten equation of enzyme activity to extract relevant parameters (fitted line in red).

**Fig S6:**
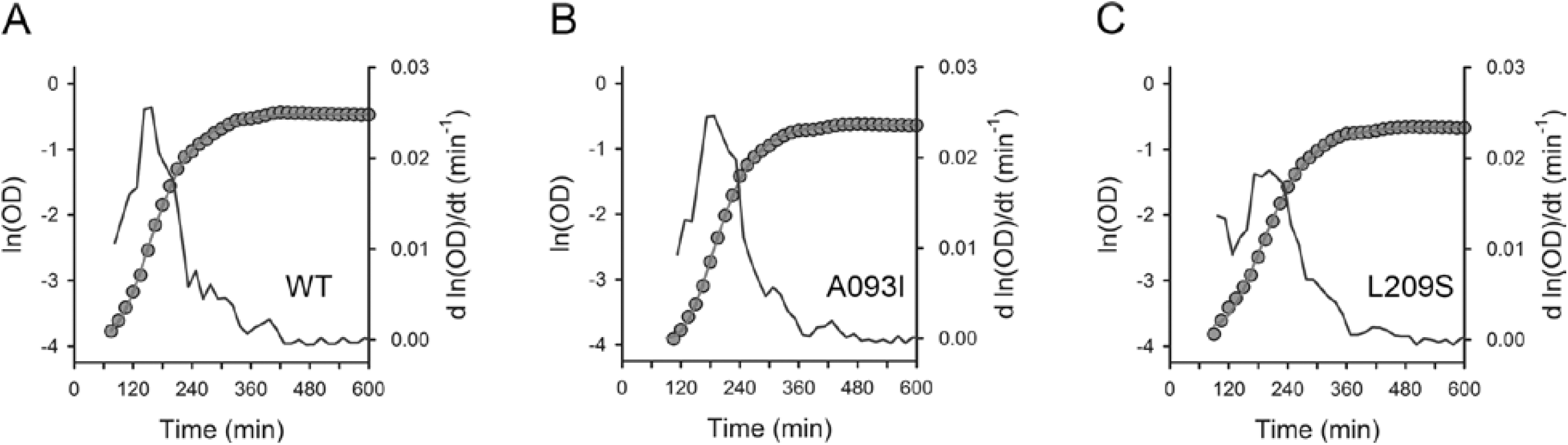
Representative growth curves of (A) WT, (B) A093I, and (C) L209S strains. Each growth curve is shown as ln(OD) vs time plot (left y-axis). The experimental data is shown in gray circles and the Gompertz fit is shown in solid red line. The instantaneous time derivative of the ln(OD) data is shown in blue line (right y-axis). The strains were chosen to illustrate the quality of the fit across different range of growth rates and lag times (see Table S2 for growth parameters).

**Fig S7:**
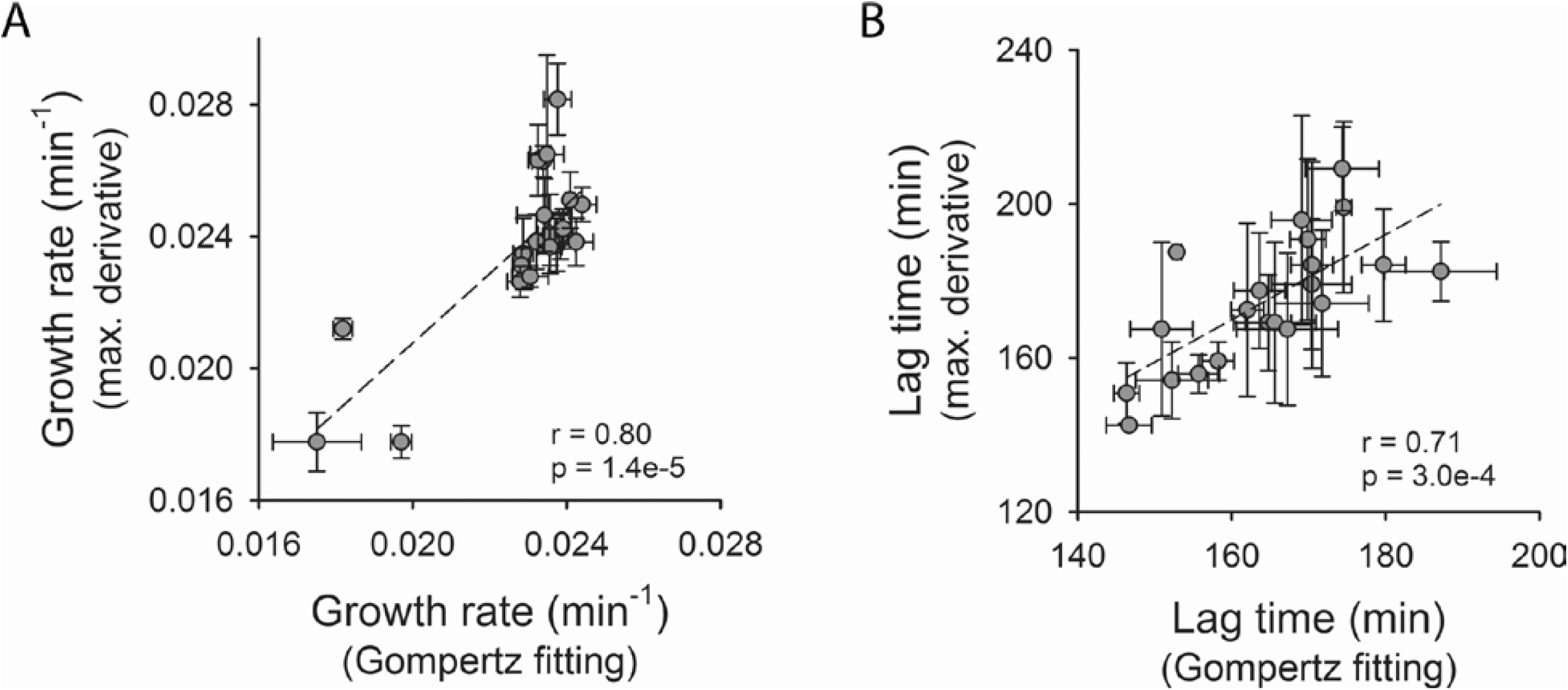
Correlation between growth parameters derived from Gompertz fitting and maximum-derivative method. The parameters derived from both the methods correlate very well as indicated by Pearson’s correlation parameters (r and p-values). The data points represent mean and error bars are standard deviation of 6 or 9 measurements (see Table S2).

**Fig S8:**
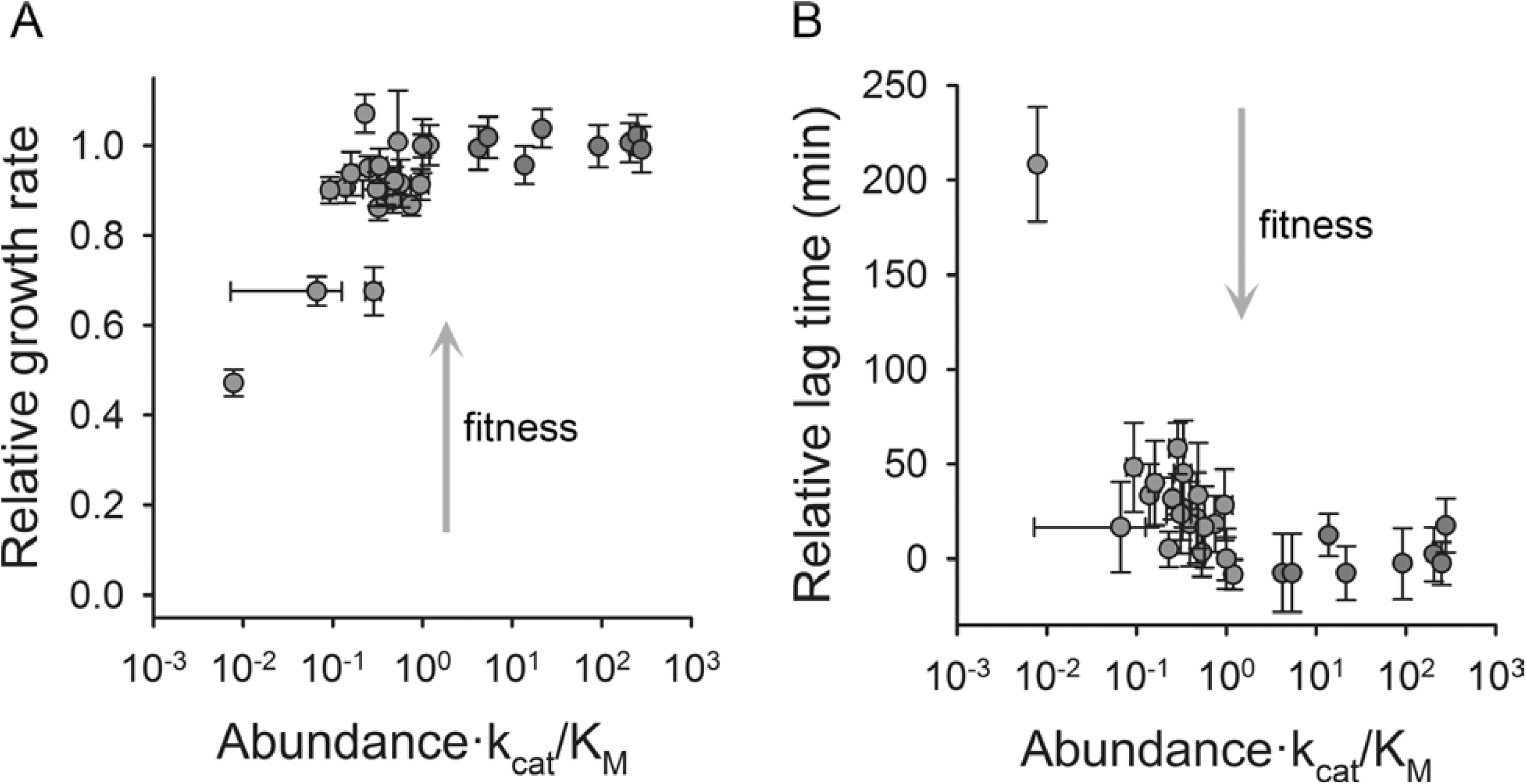
Traits of population growth. (A) Relative growth rates(*μ*/*μ_*WT*_*) and (B) relative lag time (*λ* − *λ_*WT*_*) obtained from analysis of growth curve derivatives shown as a function of catalytic capacity which is defined as abundance × *k*_*cat*_/*K*_*M*_. The mutant data is shown in gray circles, whereas red circles represent the BW27783 strain with varying degrees of overexpression of WT Adk from a pBAD plasmid. Data for WT is shown in green. Fig 2 is an equivalent figure with growth rate and lag times obtained after fitting the raw data with Gompertz equation (Eq. 2).

**Fig S9:**
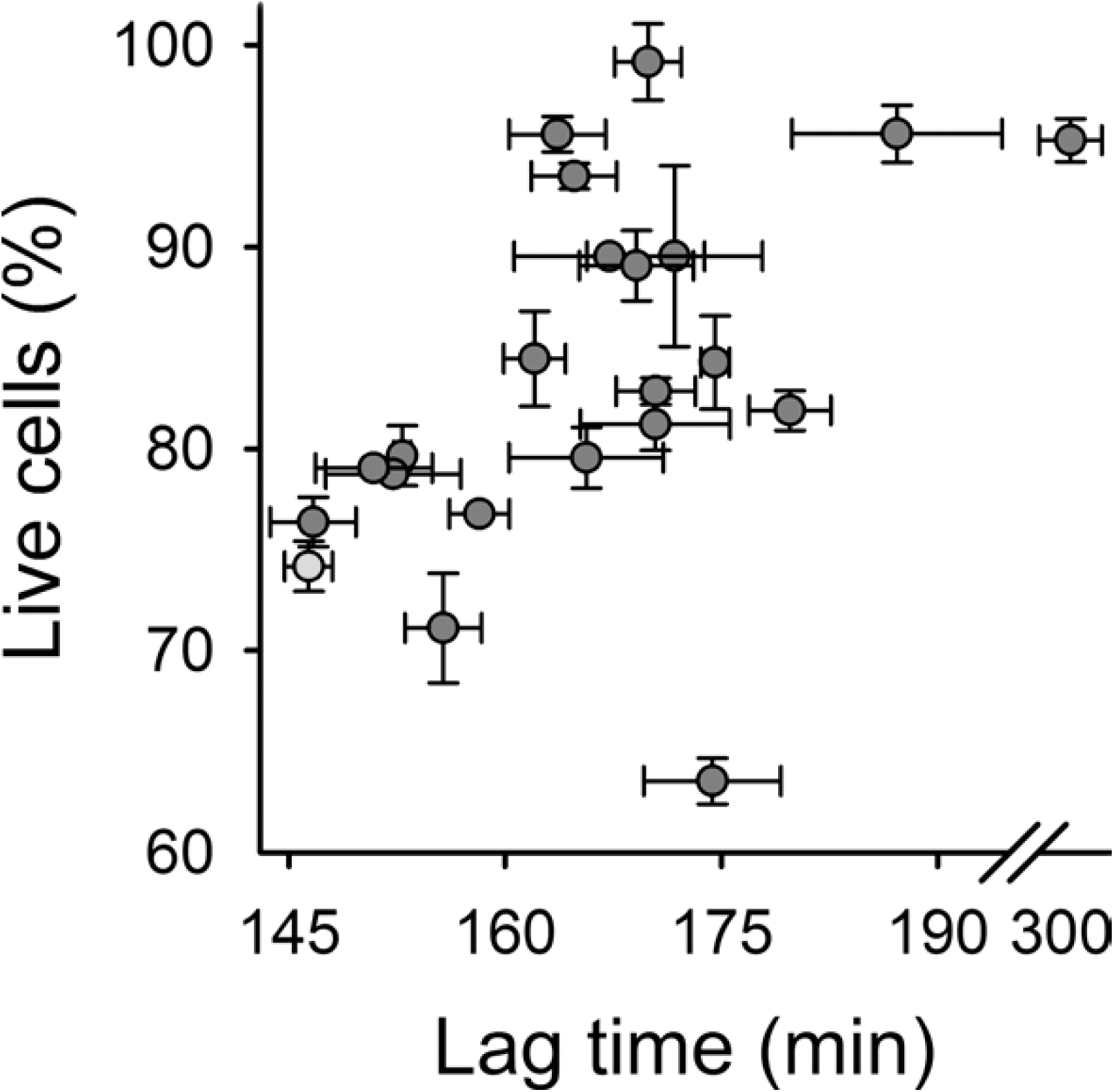
Percentage of live or viable cells of WT and mutant Adk strains at saturation (16 hours of growth) versus their population lag time. The cultures were grown overnight at 30 °C, and then stained using fluorescent dyes Syto9 (specific for live cells) and propidium iodide (specific for dead cells). The data points are mean and error bars represent standard deviation of 2 biological replicates. WT Adk strain is shown in green.

**Fig S10:**
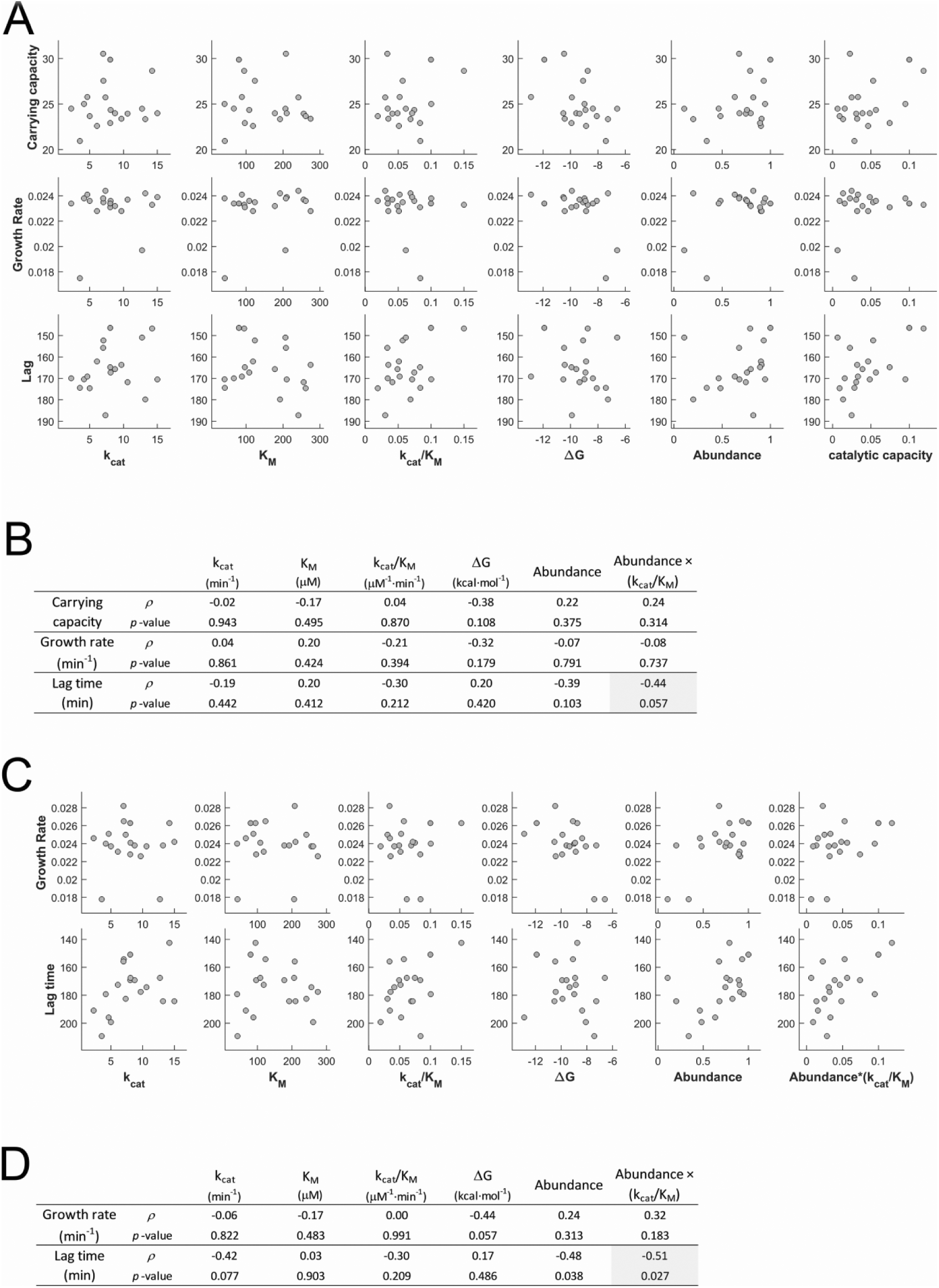
Scatter plots of growth parameters (carrying capacity, growth rate and lag times) and molecular and cellular properties of Adk. Parameters were obtained using (A) Gompertz fit and (C) analysis of growth curve derivatives. Panels (B) and (D) show Spearman’s correlation coefficients (*ρ*) and *p*-values for each of the sub-plots in panels (A) and (C) respectively. The highest correlation values in each panel are highlighted in yellow. V106N was excluded from all correlation calculations.

**Fig S11:**
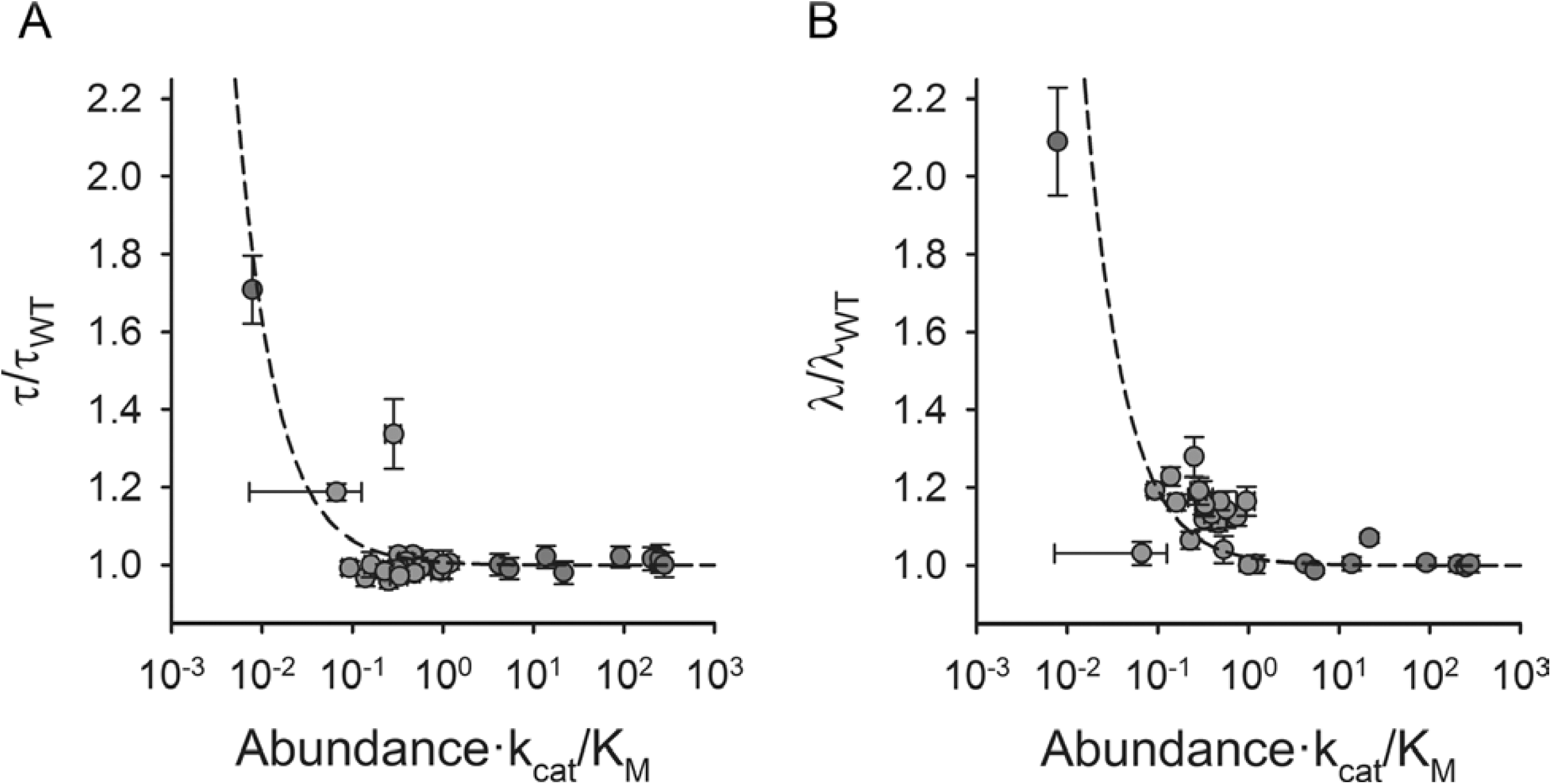
(A) Relative growth time (*τ*/*τ_*WT*_*) and (B) relative lag time (*λ*/*λ*_*WT*_) as a function of catalytic capacity (abundance × *k*_*cat*_/*K*_*M*_). The dashed line shows a fit to Eq. 3, where the asymptote (*a*) was assumed to be 1. The K_M_-like parameter *b* for growth time was 0.006 and that for lag time was 0.019, which indicates that the WT catalytic capacity is closer to the cusp for lag time than for growth time. The mutant data is shown in gray circles, whereas the overexpression data is shown in red. In green is shown WT, while the blue circle indicates V106N which was omitted from the fitting. The error bars represent standard deviation of parameters derived from growth curves of 3 colonies (biological replicates) in triplicates (9 curves). See Table S2 and S3 for the parameters.

**Fig S12:**
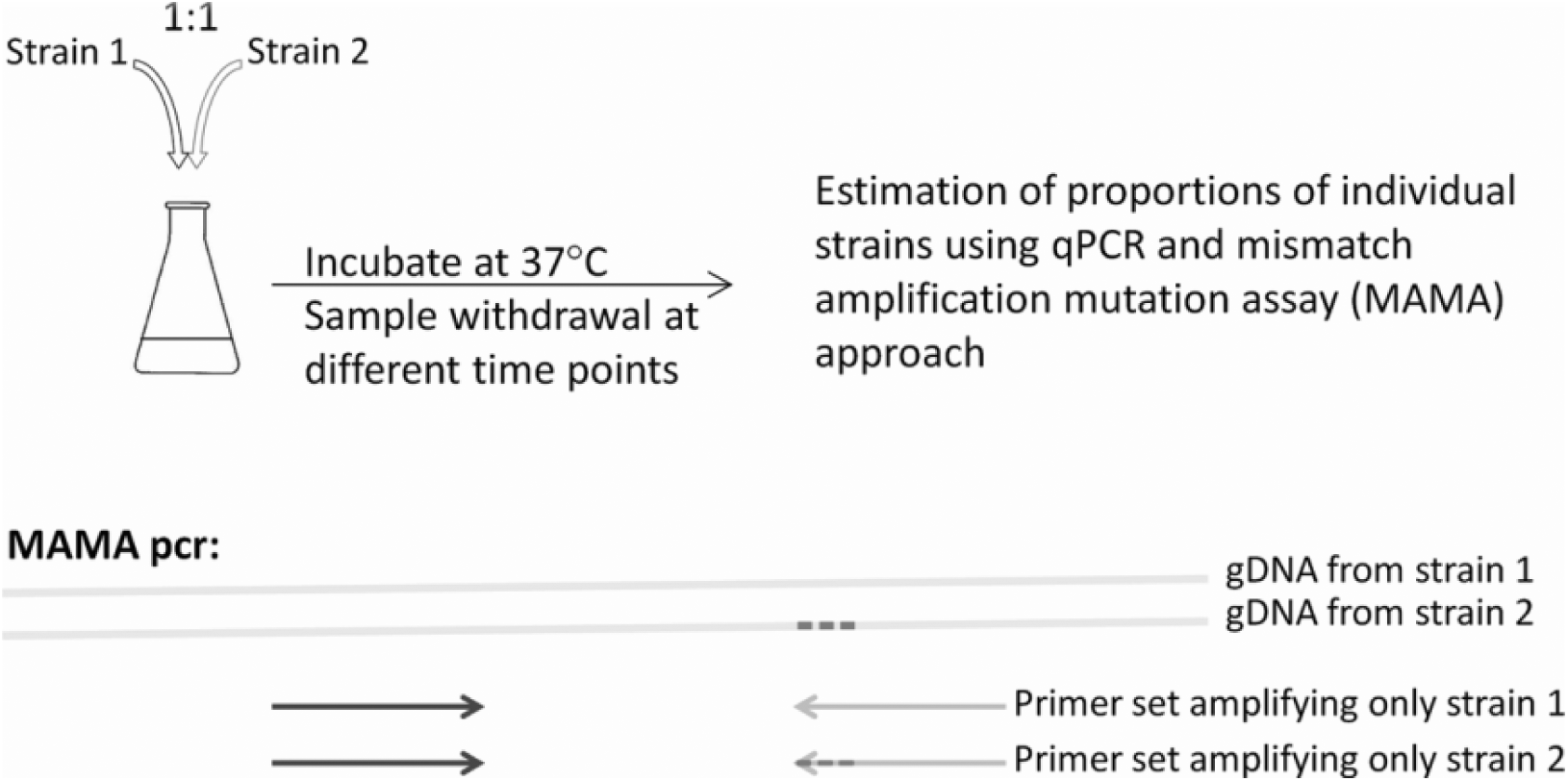
Schematic representation of binary growth competition experiments and estimation of relative proportion of competing strains. The strains (1) and (2) are mixed in 1:1 proportion and were grown at 37 °C. Samples were drawn at different time points, normalized for OD, and genomic DNA was extracted. The proportions of individual strains were estimated by a qPCR method employing mismatch amplification mutation assay method (see Methods). We designed a set of primers to differentially amplify the strains by matching the 3’-end of one of the primers to the site of mutation and using Taq DNA polymerase for amplification.

**Table S1:** Structural and biophysical parameters

| adk strain | SC acc<sup>a</sup><br>(%) | Residue depth (Å) | Fraction of<br>WT in MSA | Fraction of<br>mutant in MSA<sup>b</sup> | T<sub>m</sub> (DSC)<br>(°C) | T<sub>m</sub> (TFA)<sup>c</sup><br>(°C) | ΔG<sup>d</sup><br>(kcal/mol) | k<sub>cat</sub><sup>e</sup><br>(min<sup>-1</sup>) | K<sub>M</sub><sup>f</sup><br>(μM) | k<sub>cat</sub>/K<sub>M</sub><br>(μM<sup>-1</sup>min<sup>-1</sup>) |
| --- | --- | --- | --- | --- | --- | --- | --- | --- | --- | --- |
| WT | -- | -- | -- | -- | 55.9 | 53.8 | -11.9 | 8.05 | 80.58 | 9.99E-02 |
| L082F | 0.0 | 8.8 | 0.57 | 0.00 | 49.2 | 47.4 | -8.7 | 14.23 | 94.86 | 1.50E-01 |
| L082V | 0.0 | 8.8 | 0.57 | 0.14 | 55.7 | 53.2 | -11.3 | n.d.<sup>g</sup> | n.d.<sup>g</sup> | n.d.<sup>g</sup> |
| L083A | 0.1 | 9.0 | 0.70 | 0.00 | 48.5 | 46.9 | -8.9 | 6.10 | 118.73 | 5.13E-02 |
| L083F | 0.1 | 9.0 | 0.70 | 0.60 | n.d.<sup>g</sup> | 54.2 | -10.8 | n.d.<sup>g</sup> | n.d.<sup>g</sup> | n.d.<sup>g</sup> |
| L083I | 0.1 | 9.0 | 0.70 | 0.23 | 52.4 | 50.7 | -10.4 | 9.70 | 274.80 | 3.53E-02 |
| L083T | 0.1 | 9.0 | 0.70 | 0.00 | 49.4 | 49.3 | -9.9 | 8.05 | 96.55 | 8.34E-02 |
| A093F | 0.0 | 5.9 | 0.85 | 0.00 | 49.4 | 47.2 | -9.1 | 7.05 | 123.83 | 5.69E-02 |
| A093I | 0.0 | 5.9 | 0.85 | 0.00 | 51.3 | 49.7 | -9.9 | 7.34 | 241.75 | 3.04E-02 |
| A093L | 0.0 | 5.9 | 0.85 | 0.07 | 50.6 | 47.9 | -9.6 | 8.75 | 178.05 | 4.91E-02 |
| A093Y | 0.0 | 5.9 | 0.85 | 0.00 | 51.2 | 49.4 | -9.3 | 10.61 | 256.23 | 4.14E-02 |
| V106A | 0.0 | 7.4 | 0.74 | 0.38 | 49.7 | 47.3 | -9.0 | 4.19 | 41.78 | 1.00E-01 |
| V106H | 0.0 | 7.4 | 0.74 | 0.00 | 43.2 | 41.2 | -6.6 | 12.74 | 206.54 | 6.17E-02 |
| V106L | 0.0 | 7.4 | 0.74 | 0.04 | 50.1 | 48.5 | -8.9 | 8.10 | 108.60 | 7.46E-02 |
| V106N | 0.0 | 7.4 | 0.74 | 0.00 | 39.0 | 37.9 | -4.8 | 12.07 | 158.06 | 7.64E-02 |
| V106W | 0.0 | 7.4 | 0.74 | 0.04 | 45.0 | 43.0 | -7.3 | 13.22 | 192.05 | 6.88E-02 |
| Y182F | 7.2 | 6.3 | 0.86 | 0.14 | 55.4 | 53.8 | -10.5 | 15.00 | 210.25 | 7.13E-02 |
| Y182V | 7.2 | 6.3 | 0.86 | 0.00 | 45.7 | 46.2 | -8.1 | 5.01 | 261.53 | 1.92E-02 |
| L209A | 0.0 | 7.0 | 0.23 | 0.08 | 45.0 | 44.3 | -8.4 | 2.30 | 66.94 | 3.43E-02 |
| L209F | 0.0 | 7.0 | 0.23 | 0.01 | 51.5 | 51.0 | -10.5 | 7.00 | 208.17 | 3.36E-02 |
| L209I<sup>h</sup> | 0.0 | 7.0 | 0.23 | 0.58 | 56.1 | 55.5 | -12.9 | 4.66 | 88.82 | 5.25E-02 |
| L209S | 0.0 | 7.0 | 0.23 | 0.00 | 43.2 | 42.4 | -7.4 | 3.57 | 42.55 | 8.38E-02 |

<sup>a</sup> % sidechain accessibility calculated using coordinates of pdb 4ake<sup>b</sup> fraction in multiple sequence alignment when WT amino acid is excluded<sup>c</sup> melting temperature from thermofluor assay<sup>d</sup> derived from isothermal urea denaturation experiments at 25C<sup>e</sup> kcat for ADP formation<sup>f</sup> K<sub>M</sub> for ATP<sup>g</sup> not determined<sup>h</sup> the only case in this dataset where fraction of mutant amino acid was greater than WT amino acid in MSA

**Table S2:** Intracellular abundance and growth parameters of adk mutants

| adk strain | Carrying capacity, $K^a$ | s.d. in $K^b$ | Growth rate, $\mu^a$ (min<sup>-1</sup>) | s.d. in $\mu^b$ | Lag time, $\lambda^{a,c}$ (min) | s.d. in $\lambda^b$ | Abundance<sup>d</sup> | s.d. in abundance<sup>e</sup> |
| --- | --- | --- | --- | --- | --- | --- | --- | --- |
| WT | 29.9 | 3.0 | 0.0234 | 2.85E-04 | 146.4 | 1.7 | 1.00 | 0.00 |
| L082F | 28.7 | 2.9 | 0.0233 | 2.40E-04 | 146.7 | 3.0 | 0.79 | 0.05 |
| L082V | 24.4 | 0.8 | 0.0229 | 2.60E-04 | 158.2 | 2.1 | n.d.<sup>f</sup> | n.d.<sup>f</sup> |
| L083A | 22.6 | 0.8 | 0.0228 | 2.22E-04 | 162.0 | 2.1 | 0.90 | 0.08 |
| L083F<sup>g</sup> | 27.3 | 1.3 | 0.0182 | 2.48E-04 | 152.9 | 0.6 | 1.19 | 0.10 |
| L083I | 23.4 | 0.9 | 0.0228 | 3.30E-04 | 163.6 | 3.3 | 0.91 | 0.02 |
| L083T | 22.9 | 1.0 | 0.0231 | 4.72E-04 | 164.8 | 2.9 | 0.89 | 0.03 |
| A093F | 27.6 | 4.3 | 0.0235 | 4.37E-04 | 152.2 | 4.7 | 0.93 | 0.03 |
| A093I<sup>g</sup> | 25.7 | 0.5 | 0.0244 | 3.69E-04 | 187.2 | 7.3 | 0.82 | 0.07 |
| A093L | 24.0 | 1.0 | 0.0232 | 3.44E-04 | 165.6 | 5.3 | 0.80 | 0.13 |
| A093Y | 24.0 | 1.1 | 0.0237 | 4.90E-04 | 171.7 | 6.1 | 0.75 | 0.23 |
| V106A | 25.0 | 0.5 | 0.0238 | 3.28E-04 | 170.4 | 5.2 | 0.95 | 0.21 |
| V106H<sup>g</sup> | 24.5 | 2.2 | 0.0197 | 2.68E-04 | 150.9 | 4.1 | 0.11 | 0.10 |
| V106L | 24.4 | 2.2 | 0.0236 | 3.08E-04 | 167.2 | 6.6 | 0.76 | 0.05 |
| V106N | 5.9 | 0.3 | 0.0137 | 6.85E-04 | 305.8 | 20.0 | 0.01 | 0.00 |
| V106W | 23.4 | 1.0 | 0.0242 | 4.48E-04 | 179.7 | 2.8 | 0.20 | 0.01 |
| Y182F | 24.0 | 1.3 | 0.0239 | 2.15E-04 | 170.4 | 2.8 | 0.68 | 0.09 |
| Y182V | 23.7 | 1.1 | 0.0236 | 3.08E-04 | 174.6 | 1.0 | 0.48 | 0.08 |
| L209A | 24.5 | 1.3 | 0.0234 | 7.12E-04 | 169.9 | 2.3 | 0.46 | 0.07 |
| L209F | 30.5 | 1.8 | 0.0238 | 3.54E-04 | 155.7 | 2.7 | 0.68 | 0.05 |
| L209I | 25.8 | 1.2 | 0.0241 | 1.27E-04 | 169.1 | 4.0 | 0.63 | 0.14 |
| L209S | 20.9 | 1.5 | 0.0175 | 1.15E-03 | 174.4 | 4.8 | 0.34 | 0.07 |

<sup>a</sup> parameters derived by fitting Gompertz equation to ln(OD) vs time at 37 C
<sup>b</sup> standard deviation derived from 9 replicates (triplicates of 3 biological replicates)
<sup>c</sup> time required to achieve maximum growth rate
<sup>d</sup> abundance measured after 4h of growth at 37 C
<sup>e</sup> standard deviation derived 2 biological replicates
<sup>f</sup> not determined
<sup>g</sup> growth parameters derived for 6 replicates (triplicates of 2 biological replicates)

**Table S3:** Intracellular abundance and growth parameters of WT adk overexpression from pBAD plasmid in *E. coli* BW27783 strain

| arabinose<br>concentration<br>(%) | Carrying capacity,<br>$K^a$ | s.d. in $K^b$ | Growth rate, $\mu^a$<br>($\text{min}^{-1}$) | s.d. in $\mu^b$ | Lag time, $\lambda^{a,c}$<br>(min) | s.d. in $\lambda^b$ | Abundance<sup>d</sup> | s.d. in abundance<sup>e</sup> |
| --- | --- | --- | --- | --- | --- | --- | --- | --- |
| no plasmid | 29.5 | 2.5 | 0.0195 | 5.10E-04 | 126.7 | 1.0 | 1.00 | 0.10 |
| 0.00E+00 | 32.0 | 0.5 | 0.0195 | 2.00E-04 | 127.2 | 0.7 | 4.20 | 0.42 |
| 3.05E-06 | 30.9 | 1.8 | 0.0191 | 2.00E-04 | 127.2 | 2.0 | 13.73 | 1.37 |
| 1.22E-05 | 31.3 | 1.0 | 0.0197 | 2.00E-04 | 125.0 | 0.4 | 5.39 | 0.54 |
| 4.88E-05 | 27.1 | 1.0 | 0.0199 | 2.65E-04 | 135.7 | 1.1 | 21.55 | 2.15 |
| 1.95E-04 | 32.7 | 0.9 | 0.0191 | 1.00E-04 | 127.5 | 1.2 | 91.27 | 9.12 |
| 7.81E-04 | 31.6 | 2.5 | 0.0192 | 2.52E-04 | 127.0 | 2.1 | 205.31 | 20.51 |
| 3.13E-03 | 32.6 | 1.6 | 0.0192 | 4.73E-04 | 126.1 | 1.4 | 250.39 | 25.02 |
| 5.00E-02 | 28.4 | 3.1 | 0.0195 | 3.61E-04 | 127.1 | 2.5 | 278.69 | 27.84 |

<sup>a</sup> parameters derived by fitting Gompertz equation to $\ln(\text{OD})$ vs time at 37 C
<sup>b</sup> standard deviation of 3 replicates
<sup>c</sup> time required to achieve maximum growth rate
<sup>d</sup> abundance measured after 4h of growth at 37 C
<sup>e</sup> standard deviation of 2 biological replicates

